# CLUH interactome reveals an association to SPAG5 and a proximity to the translation of mitochondrial protein

**DOI:** 10.1101/2021.07.08.451585

**Authors:** Mickaële Hémono, Alexandre Haller, Johana Chicher, Anne-Marie Duchêne, Richard Patryk Ngondo

## Abstract

Mitochondria require thousands of proteins to fulfil their essential function in energy production and other fundamental biological processes. These proteins are mostly encoded by the nuclear genome, translated in the cytoplasm before being imported into the organelle. RNA binding proteins (RBPs) are central players in the regulation of this process by affecting mRNA translation, stability or localization. CLUH is an RBP recognizing specifically mRNAs coding for mitochondrial proteins, but its precise molecular function and interacting partners remain undiscovered in mammals. Here we reveal for the first time CLUH interactome in mammalian cells. Using both co-IP and BioID proximity-labeling approaches, we identify novel molecular partners interacting stably or transiently with CLUH in HCT116 cells and mouse embryonic stem cells. We reveal a stable RNA-independent interaction of CLUH with itself and with SPAG5 in cytosolic granular structures. More importantly, we uncover an unexpected proximity of CLUH to mitochondrial proteins and their cognate mRNAs in the cytosol. Additionally, our data highlight the importance of CLUH TPR domain for its interactions with both proteins and mRNAs. Overall, through the analysis of CLUH interactome, our study sheds a new light on CLUH molecular function by highlighting its association to the translation and subcellular localization of some mRNAs coding for mitochondrial proteins.

## INTRODUCTION

Mitochondria are considered as the “powerhouses of the cell” as they generate ATP through oxidative phosphorylation and are involved in many other cellular functions, such as autophagy, apoptosis and metabolic homeostasis [1]. They are semi-autonomous organelles because most of the essential mitochondrial proteins are synthesized by cytoplasmic ribosomes and then imported. The mammalian mitochondrial proteome is composed of more than 1100 nuclear encoded proteins while only 13 are encoded by the mitochondrial genome [2]. The principal pathway to import mitochondrial proteins, consists in a complete cytosolic translation, followed by chaperone-mediated transfer to the outer mitochondrial membrane (OMM) and a translocation through the TOM complex [3]. Most of the mitochondria-destined precursor proteins contain a N-terminal mitochondrial targeting sequence (MTS) that is cleaved during the import to release the mature protein. An alternative pathway suggests that translation in the vicinity of the OMM is important for efficient mitochondrial targeting [3, 4]. This co-translational import mechanism has been largely studied in yeast [5] but evidence in mammals is generally missing. A very recent study revealed several hundred mRNAs coding for mitochondrial proteins enriched at the OMM in mammalian cells in a translation dependent manner, supporting the idea of OMM localized translation [6]. The localization of mRNAs at the OMM has also been demonstrated in other higher eukaryotes such as plants [7] and insects [8].

Regardless the protein-import pathway, mRNAs coding for mitochondrial proteins need to be distinguished from other cellular mRNAs and handled specifically. This is mainly achieved by RNA binding proteins (RBPs) that can regulate multiple aspects of mRNA life cycle, affecting its stability or degradation, localization and translation efficiency [9]. Several RBPs are involved in regulating mitochondrial functions, but most of them have been identified in yeast and are not always functionally conserved in mammals [5, 10].

CLUH (clustered mitochondria homolog) is a RNA binding protein that has been found associated with transcripts coding for mitochondrial proteins in mammals [11] and described as promoting translation and stability of at least a subset of these [12]. *CLUH* deletion in cultured mammalian cell lines causes mitochondrial clustering around the nucleus, suggesting its involvement in mitochondrial biogenesis through a mechanism that remains still elusive (Gao et al., 2014; Pla-Martin et al., 2020; Wakim et al., 2017). This abnormal mitochondrial phenotype has also been observed upon deletion of *CLUH* orthologs in plants [13], *Drosophila melanogaster* [14], *Dictyostelium* [15] and yeast [16]. In mice, CLUH deficiency causes neonatal lethality [12], presumably due to a drastic oxidative phosphorylation (OXPHOS) and metabolic defects [17]. In fact, after birth, the energy demand of some tissues is considerably increased and coincides with a metabolic switch from a major anaerobic glycolytic toward a mainly aerobic OXPHOS metabolism [18]. Such a metabolic switch also occurs during the process of differentiation of pluripotent stem cells toward somatic cells and is accompanied by a maturation of mitochondria with a remarkable change of morphology and localization [19].

To our knowledge, despite a description of the interaction of CLUH ortholog with the ribosome at the mitochondrial surface in *Drosophila* [20], nothing is known about the molecular partners of CLUH. Even though CLUH has been described to have a crucial role for mitochondrial function in the liver and to affect mitochondrial mRNA translation [21], stability [12] and localization [22], its precise molecular function and associated protein interactors remain to be discovered.

In the present work, we took advantage of two complementary proteomic approaches to identify both stable and transient CLUH interactions in mammalian cultured cells. First, we identified by co-immunoprecipitation, novel stable interactions of CLUH with itself and with the SPAG5/KNSTRN complex. We showed that these interactions are RNA independent and require the TPR protein-protein interaction domain of CLUH. Using a split-sfGFP [23] approach we found that CLUH interacts with SPAG5 in cytosolic granular structures. Afterward, using both BioID2 [24] and TurboID [25] proximity labeling approaches, we identified for the first time <u>C</u>LUH <u>P</u>roximal <u>M</u>itochondrial <u>P</u>roteins (CPMPs). We found that CLUH interact transiently with those nuclear encoded mitochondrial proteins before their import into the organelle. Moreover, we found that CLUH interacts also with the mRNAs coding for those CPMPs, with no evident impact on their cytosolic translation. Furthermore, our data point toward a role of CLUH in the subcellular localization of mRNAs coding for some CPMPs. Taken together, our data shed a new light on to CLUH molecular function, by revealing for the first time its interactors and its physical proximity to both the mRNA and the resultant translated mitochondrial proteins.

## RESULTS

### Identification of CLUH interacting proteins in both human HCT116 and mESCs

In order to better understand CLUH functions we analyzed its interactome by co-immunoprecipitation (Co-IP) followed by Liquid Chromatography with tandem mass spectrometry (LC-MS/MS) in both human HCT116 cells and mouse embryonic stem cells (mESCs) (Figure 1A and 1B). We generated a polyclonal HCT116 stable cell line ectopically expressing mouse CLUH (mCLUH) protein tagged with a triple HA epitope (3xHA) (Figure S1 A). We also engineered mESCs, using CRISPR/cas9 genome editing approach, in order to express an endogenously 3xHA-tagged CLUH protein (Figure S1B and S1C). Co-IP experiments, using anti-HA-coupled beads, were performed on both cell types expressing 3xHA-CLUH (IP CLUH) and on wild-type cells (IP mock) used as non-specific control (Figure S1D). Three replicate experiments per condition were analyzed by LC-MS/MS : more than 500 proteins were identified per condition and quantified using label-free spectral counting method (Figure 1C, 1D, Table S1 and S2) [26]. Significant enrichment of the IP CLUH versus IP mock control was defined by a threshold a fold change (FC) of 2 and 10% false discovery rate (FDR). In HCT116 cells, only 5 significantly enriched proteins were identified (in red, Figure 1E) among which CLUH is the most enriched followed by two mitotic spindle associated proteins SPAG5 and KNSTRN. The two proteins, also known as Astrin and Kinastrin/SKAP, form a complex that localizes to microtubules plus ends [27]. On the other hand, in mESCs we identified 55 significantly enriched proteins (Figure 1F), largely corresponding to RNA binding proteins (Figure S1E). The only identified CLUH-interacting proteins common to both HCT116 and mESC are SPAG5 and KNSTRN.

**Figure 1:**
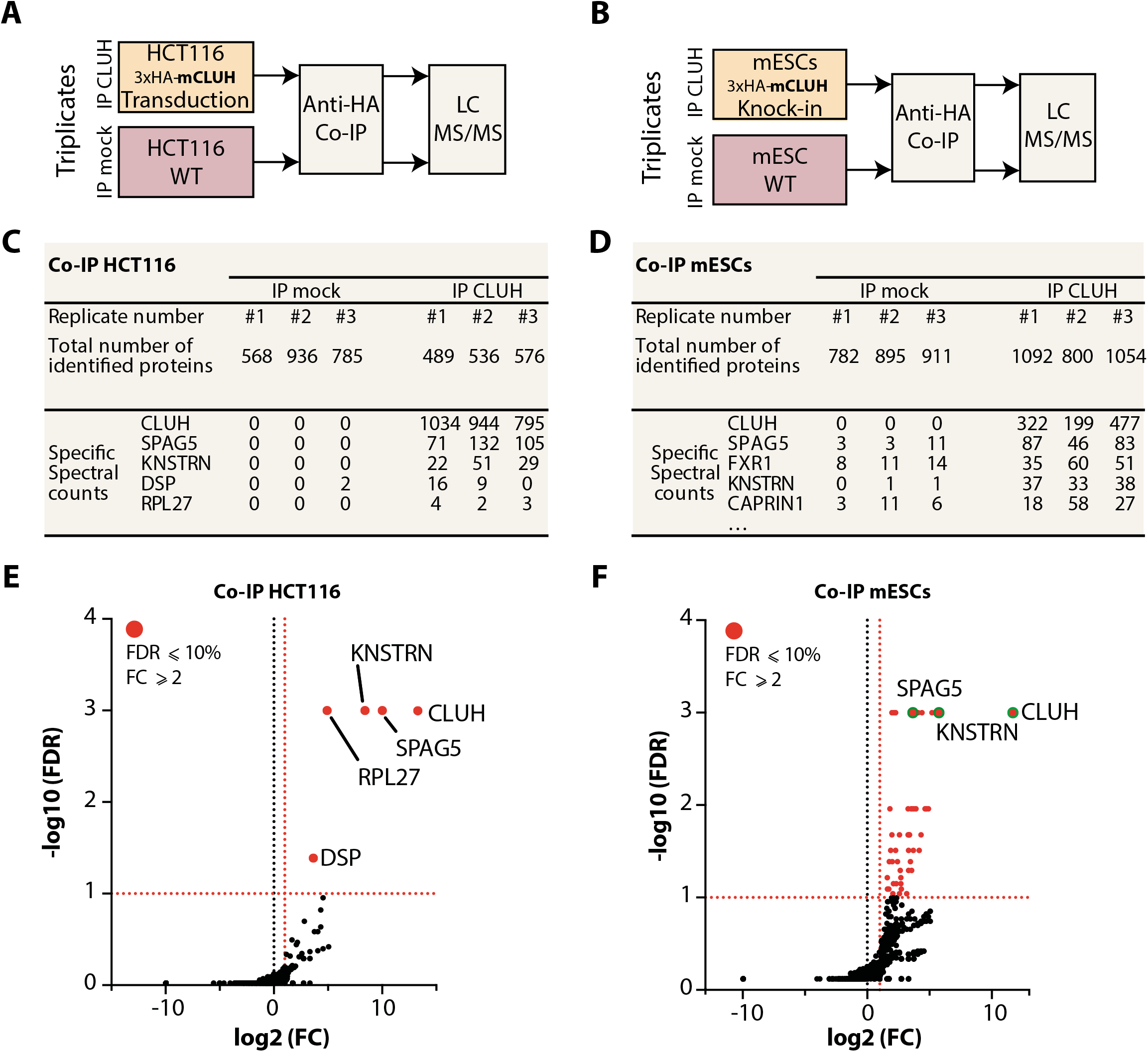
Identification of CLUH interractome by co-immunoprecipitation in both HCT116 cells and mESCs. **(A-B)** Schematic representation of the co-IP experimental design. Co-immunoprecipitated proteins from 3xHA-mCLUH sample (IP CLUH) and control sample (IP mock) are identified LC MS/MS. **(A)** HCT116 cells are transduced to express 3xHA-mCLUH protein. **(B)** mESC are genome-edited to express an endogenous 3xHA-CLUH protein. The mESCs knock-in clone G12 is used (See Figure S1C). **(C-D)** Tables summarizing the MS protein identification in HCT116 **(C)** and mESCs **(D)**. Total number of proteins identified by mascot with a false discovery rate (FDR) below 1% in IP mock and IP CLUH samples. Five proteins with the highest specific spectral counts in the IP CLUH are shown. Biological replicate samples are numbered from #1 to #3. **(E-F)** Volcano plots showing the global enrichment of proteins in IP CLUH versus the IP mock. The x-axis shows the log2 fold change (FC) and the y-axis shows the −log10 of the FDR (n=3), obtained using SAINTexpress software [26]. Significantly enriched proteins are shown in red and are defined by a fold change greater than two and a FDR < 0.1 (shown as dashed red line). Selected proteins with the highest spectral count (shown in D) are labeled and identified with a green circle.

### CLUH interacts with both SPAG5 and KNSTRN in RNA-independent manner

To validate the interaction of endogenous human CLUH protein with the two top identified proteins, SPAG5 and KNSTRN, we generated polyclonal HCT116 stable cell lines expressing GFP-tagged bait proteins and performed co-IPs using anti-GFP coupled beads (Figure 2A). The endogenous CLUH protein co-immunoprecipitated with both GFP-KNSTRN and GFP-SPAG5. These interactions were further validated by pulling down the two GFP-tagged proteins using ectopically expressed 3xHA-CLUH protein (Figure S2A and S2B). In addition, these interactions are maintained when protein extracts are treated with RNases prior pulldown (Figure S2C, S2A and S2B), showing that CLUH interacts with both SPAG5 and KNSTRN in an RNA-independent manner.

**Figure 2:**
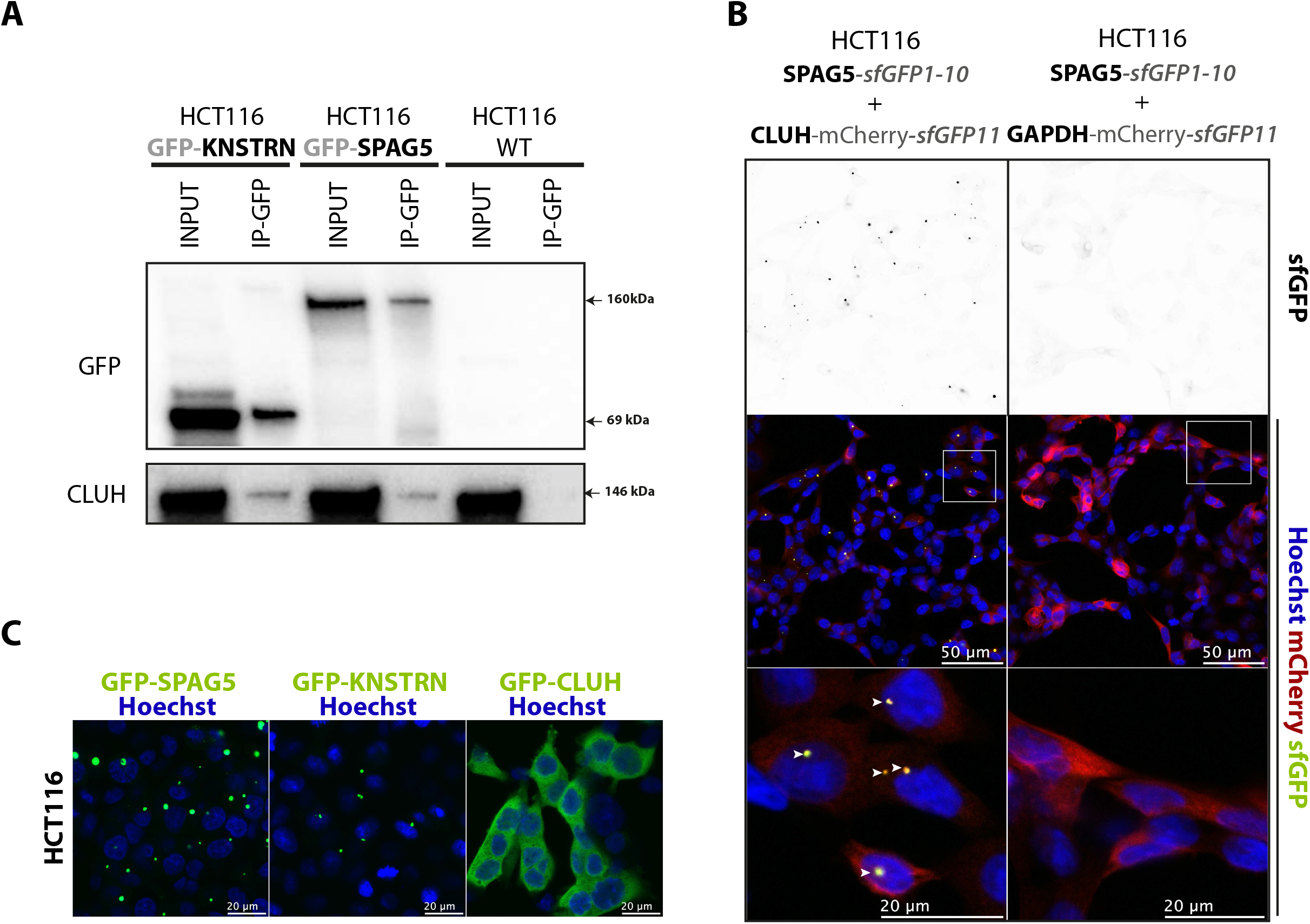
Endogenous CLUH interacts with both SPAG5 and KNSTRN and the CLUH-SPAG5 interaction occurs in granular structures. **(A)** Western blot analysis of co-IP between GFP-tagged KNSTRN or SPAG5 proteins and the endogenous CLUH protein and in HCT116 cells. The co-IP is performed using magnetic beads coupled with anti-GFP antibodies (IP-GFP) on total extracts (INPUT) from cells stably expressing the tagged proteins. **(B)** Split-GFP analysis of SPAG5 and CLUH interaction. HCT116 stably expressing SPAG5 fused with sfGFP-1-10 together with CLUH or GAPDH fused with mCherry-sfGFP11 are fixed and analyzed by confocal fluorescent microscopy. Reconstituted sfGFP signal is shown in black (upper panel) and in green (lower panels). The mCherry signal is shown in red and nuclei, stained with Hoechst, are in blue. The colocalization of green and red signal is shown in yellow and indicated with arrows. Scale bar is indicated in white. **(C)** Confocal fluorescent microscopy images of HCT116 cells stably expressing GFP fused to SPAG5, KNSTRN or CLUH. GFP signal is shown in green and nuclei, stained with Hoechst, in blue. Scale bar is indicated in white.

### CLUH interacts with SPAG5 in cytosolic granular structures

SPAG5 and KNSTRN being part of the same complex, we focused our study on SPAG5. We analyzed the CLUH-SPAG5 interaction by fluorescence microscopy using a split-sfGFP approach [23]. We selected HCT116 stable cell lines expressing SPAG5 in fusion with the truncated non-fluorescent sfGFP1-10 fragment together with CLUH or GAPDH tagged with the missing eleventh ß-strand sfGFP11 (Figure 2B). The expression of CLUH and GAPDH was followed thanks to a mCherry-fusion marker. The reconstitution of the fluorescent-competent sfGFP was only possible in the presence of CLUH and not with GAPDH control protein. Green spots, corresponding to sites of CLUH-SPAG5 interaction, were observed in perinuclear areas. Only one or two sfGFP-spots were observed per cell (Figure S2E, normal). Strikingly, this accumulation was very similar to SPAG5 and KNSTRN subcellular localization observed in HCT116 cells expressing GFP-tagged proteins (Figure 2C). Unlike CLUH that shows a globally uniform cytosolic localization (Figure 2B and 2C), SPAG5 and KNSTRN preferentially accumulated in perinuclear spot-like structures similar to previously described SPAG5 mitotic spindle localization [28, 29]. CLUH was previously described to form granules in stress condition [21], we therefore also examined CLUH localization and CLUH-SPAG5 formed structures under stress conditions (Figure S3A and S3B). Both mCherry-tagged CLUH localization and the amount of CLUH-SPAG5 formed structures did not change upon starvation or oxidative stress and were not affected by cycloheximide treatment known to disassemble stress granules. Moreover, CLUH-SPGA5 structures did not colocalize with sodium arsenite induced G3BP3-positive stress granules (Figure S3C). Overall our data indicate that CLUH interacts with SPAG5 in cytosolic structures that are not stress-granules and resembling to SPAG5/KNSTRN accumulation sites in mitotic spindle.

### CLUH self-interaction and CLUH-SPAG5 interaction require CLUH TPR domain

Given the abundance of CLUH in our co-IP data compared to prey proteins, we wondered whether it might form multimers. To verify CLUH dimerization potential, we transduced HCT116 cells to express both GFP-tagged and 3xHA-tagged CLUH and performed anti-HA co-IP followed by a western blot using antibodies against the two tags (Figure 3A). The GFP-tagged CLUH co-immunoprecipitated with 3xHA-CLUH showing that the two proteins interact with each other. This interaction is RNA-independent as it was not affected by RNase treatment prior immunoprecipitation (Figure 3B and 3A).

**Figure 3:**
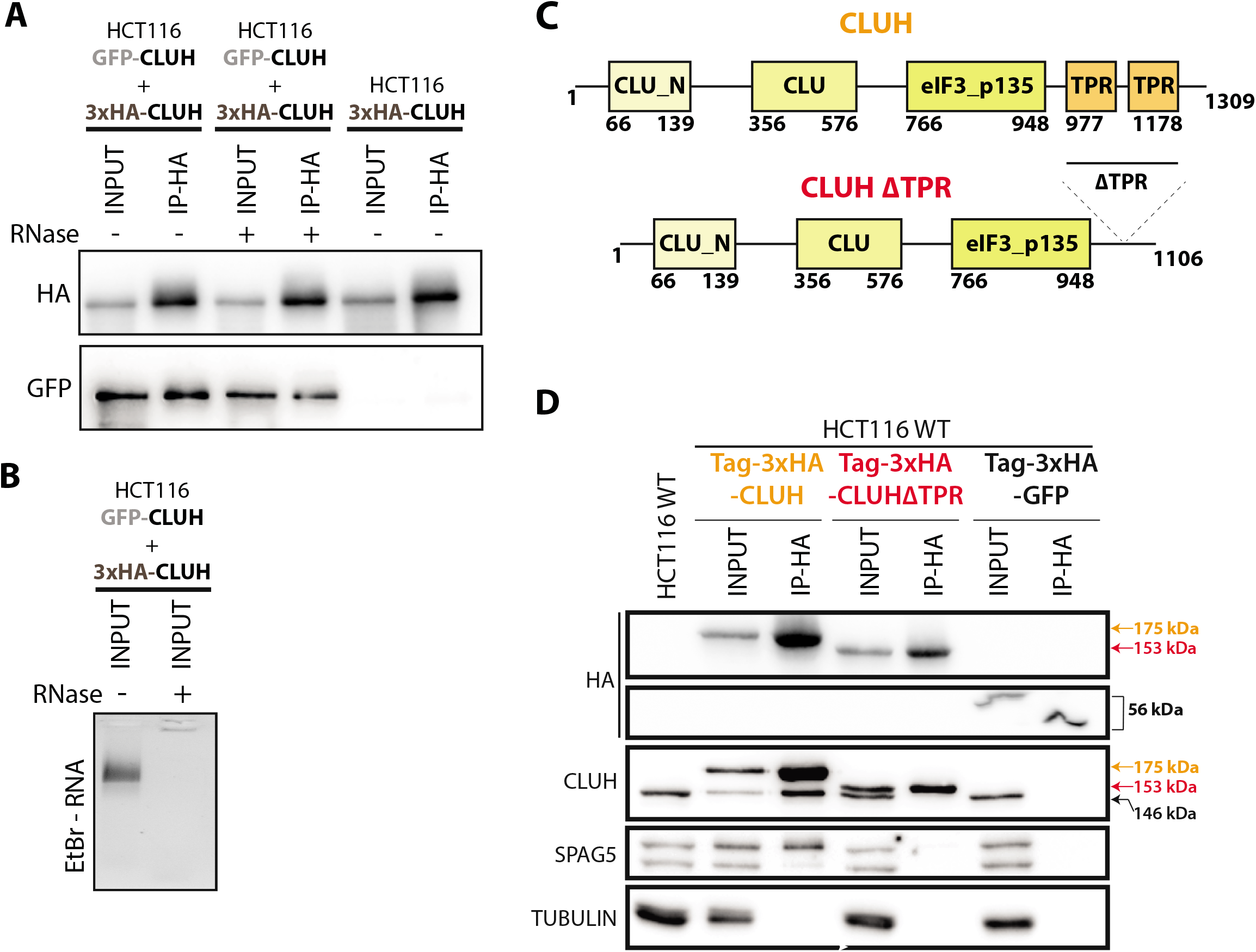
The TPR domain facilitates CLUH self-interaction and its interaction with SPAG5. **(A)** Western blot analysis of co-IP, between GFP-and 3xHA-tagged CLUH proteins stably expressed in HCT116 cells. The IP is performed using magnetic beads coupled with anti-HA antibodies (IP-HA) with protein extracts treated (+) or not (-) with RNaseA/T1. The proteins are detected using anti-HA and anti-GFP antibodies. **(B)** Ethidium bromide-stained agarose gel loaded with RNase treated (+) or non-treated (-) total protein extracts used for the IP showing the presence of ribosomal RNA. **(C)** Scheme showing the human CLUH protein domains identified using Pfam database. The amino acid positions of each domain are indicated. The mutant protein CLUHΔTPR have been generated by deleting the TPR domains. **(D)** Western blot analysis of co-IP, between 3xHA-tagged wildtype CLUH (in orange) or the CLUHΔTPR mutant (in red) with the endogenous CLUH and SPAG5 proteins. A 30 kDa tag corresponding to the BioID2 protein followed by 3xHA peptide is added in N-terminal (Tag-3xHA) of each transgene. A GFP tagged construct is used as a specificity control. The co-IP is performed on total extracts (INPUT) from HCT116 cells stably expressing the different constructs using magnetic beads coupled with anti-HA antibodies (IP-HA). The size of the endogenous CLUH (black), wildtype transgene (orange) and the delta-TPR mutant (red) is indicated. The indicated proteins are revealed using specific antibodies. TUBULIN Is used as a loading control.

The TPR domain is a protein-protein interaction module facilitating the interaction of multiple proteins [30]. The TPR domain of CLUH is very well conserved among eukaryotes and previous data in *Drosophila* highlighted its importance for Clueless (CLUH ortholog) function *in vivo* [20]. To evaluate its importance for CLUH interaction with itself and with SPAG5, we mutated CLUH protein and performed co-immunoprecipitation assays. The protein was truncated from residues 977 to 1178, generating a CLUHΔTPR protein (Figure 3C). In order to discriminate the transgene from the endogenous CLUH protein (146 kDa), we added a 30 kDa BioID2-3xHA tag in N-terminal. HCT116 cells were transduced to express tagged CLUH (175 kda), CLUHΔTPR (153 kDa) or GFP (56 kDa) proteins (Figure 3D) and the resulting protein extract used for anti-HA immunoprecipitation. Both the endogenous CLUH and SPAG5 proteins co-immunoprecipitated with the full-length CLUH protein but not with the truncated CLUHΔTPR protein nor with GFP. The annotated TPR domain of CLUH is therefore required for its self-interaction and for its interaction with SPAG5. Interestingly, these two interactions are not depending on each other as knocking down SPAG5 does not affect the efficiency of endogenous CLUH co-immunoprecipitation (Figure S4).

### CLUH proximity interactome is mainly composed of CPMPs

Co-immunoprecipitation assays only catches stable protein-protein interactions that are strong enough to be preserved under chosen experimental conditions until the elution step. Considering this limitation and to get more understanding on CLUH proximal environment, we decided on using a complementary *in vivo* BioID proximity-labeling approach [31]. We generated a polyclonal HCT116 stable cell line ectopically expressing mouse CLUH (mCLUH) protein fused, in N-terminal, to a modified biotin ligase protein (BioID2-CLUH) [24]. A control stable cell line expressing a GFP-BioID2 fusion was also generated. The addition of biotin substrate into the culture medium initiated the biotinylation of endogenous proteins proximal to the baits (Figure 4A, S5A and S5B). Biotin-labelled proteins were then isolated using streptavidin-coupled beads and identified by LC-MS/MS. Three biological replicate experiment were performed, more than 1200 proteins identified in the BioID-mCLUH condition and more than 1700 proteins in GFP-BioID2 condition (Figure 4B, Table S3). The proteins were quantified using label-free spectral counting and 90 significantly enriched proteins in BioID-mCLUH versus GFP-BioID2 background sample were identified (Figure 4C). In addition to CLUH bait protein, previously identified SPAG5 and KNSTRN were also among the most enriched proteins. Functional enrichment analysis of the whole list of CLUH proximal proteins shows an evident enrichment in terms associated with mitochondria (Figure 4D). The analysis of the protein network allowed us to highlight 43 significantly enriched mitochondrial proteins, among which 40 have an Uniprot-annotated MTS (Figure 4E). All the identified <u>C</u>LUH <u>P</u>roximal <u>M</u>itochondrial <u>P</u>roteins (CPMP) are nucleus-encoded and imported into the mitochondria. Two other noticeable groups of proteins are ribosomal subunits and some proteins functionally associated with the cytoskeleton among which are SPAG5 and KNSTRN. We also identified CLUH proximal proteins in mESC by applying the same the BioID approach. Using CRISPR/Cas9 genome engineering technology, we endogenously tagged CLUH in N-terminal with BioID2 biotin ligase (Figure S5C, S5D). Considering the importance of CLUH for mitochondrial-associated energy metabolism [17] we thought to evaluate CLUH proximity interactome in both pluripotent and differentiated mESC as the differentiation process is associated with important changes in mitochondria morphology and the energy metabolism [19]. The biotin labeling was therefore performed before and after the spontaneous mESC embryoid body differentiation process (EB) (Figure S5E). We identified 27 significantly enriched proteins in undifferentiated cells and 77 proteins in EBs with an 88.8% overlap (Figure S5F and S5G). As for HCT116 cells, we observed a strong enrichment in mitochondrial proteins (Figure S5H and S5I). Altogether, we identified 17 CPMPs in undifferentiated cells of which 14 are also found in HCT116 and 28 CPMPs in differentiated cells of which 18 were also identified in HCT116. Although we identified some common CPMPs, CLUH has evidently a different transient-interactome between HCT116 cells, undifferentiated mESC and EB-differentiated mESC. Moreover, the larger number of biotinylated proteins identified in EBs could reflect a higher CLUH participation to the increased mitochondrial activity in differentiated cells compared to pluripotent cells.

**Figure 4:**
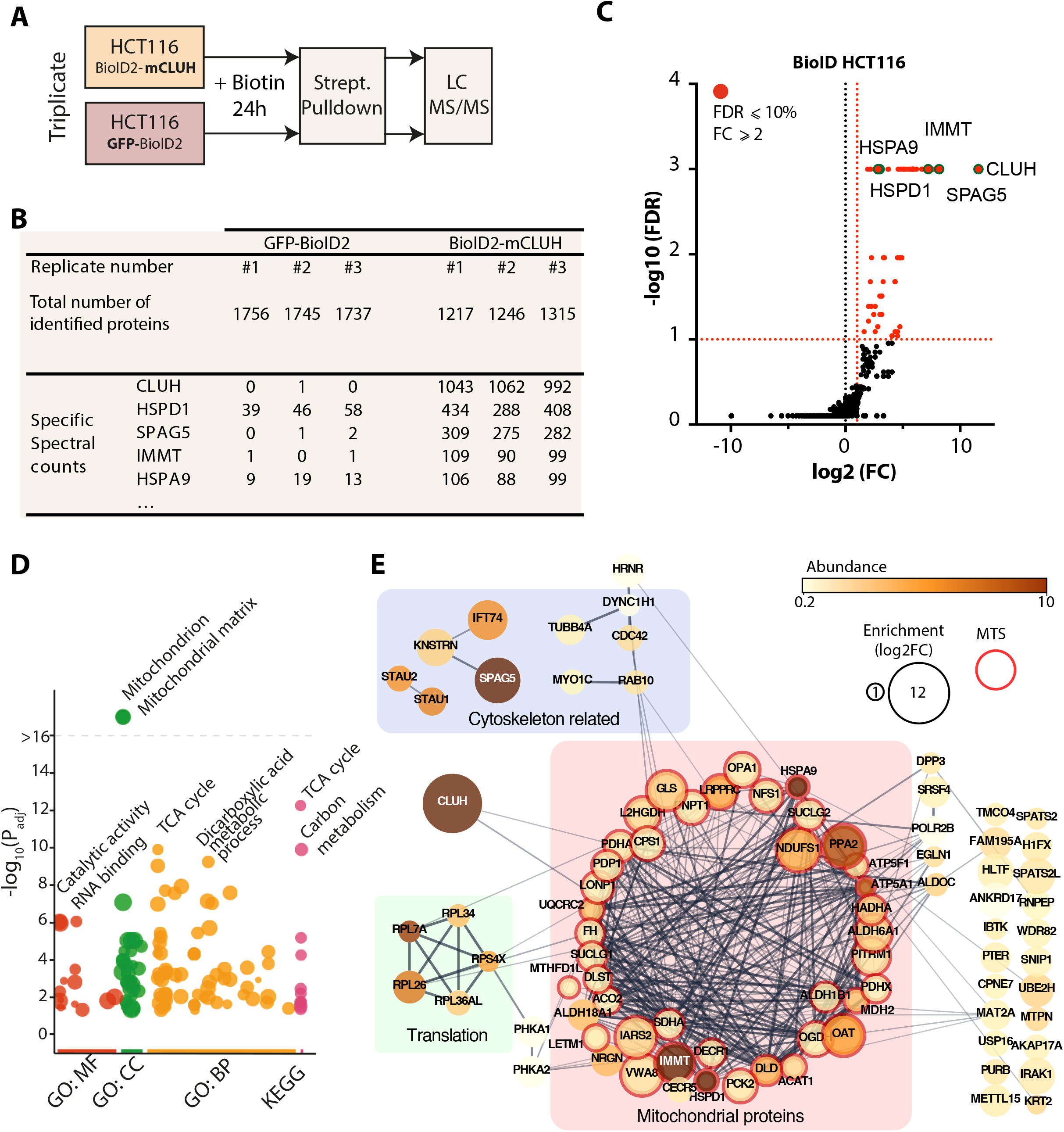
Identification of CLUH proximal proteins using BioID in HCT116 cells. **(A)** Schematic representation of the BioID experimental design using HCT116 cells stably expressing the BioID2 protein fused to the mouse CLUH (mCLUH) or GFP proteins. The proximity labeling is performed for 24 hours in the presence of 50 µM biotin in the culture medium. Biotinylated proteins, from both the specific (BioID2-CLUH) and control (GFP-BioID2) samples, are isolated using streptavidin-coupled magnetic beads and identified by LC MS/MS. **(B)** Table summarizing the MS protein identification from the BioID experiment in HCT116 cells. Total number of proteins identified by mascot with a FDR below 1%. Five proteins with the highest specific spectral counts in the BioID2-mCLUH sample are shown. Biological replicate samples are numbered from #1 to #3. **(C)** Volcano plot showing the global enrichment of proteins in BioID2-CLUH versus the GFP-BioID2 control. The x-axis shows the log2 fold change (FC), and the y-axis shows the −log10 of the false discovery rate (n=3), obtained using SAINTexpress software [26]. Significantly enriched proteins are shown in red and are defined by a fold change greater than two and a FDR < 0.1 (shown as dashed red lines). Five proteins with the highest spectral count (shown in B) are labeled and identified with a green circle. **(D)** Manhattan plot illustrating the gene ontology and pathway enrichment analysis of proteins identified in BioID experiment, generated using G:profiler tool [43]. The functional terms, associated with the protein list, are grouped in four categories: GO: MF (Molecular Function), GO: CC (Cellular Component), GO : BP (Biological Process) and KEGG pathways. The y-axis shows the adjusted enrichment p-values in negative log10 scale. The circle sizes are in accordance with the corresponding term size (in the database) and terms from the same GO subtree are located close to each other on the x-axis. The most significantly enriched terms are labeled. **(E)** Visualization of the functional interaction network of CLUH proximal proteins identified by BioID, generated with the Cytoscape StringApp [44]. The proteins have been grouped according to three most represented functional categories: “Cytoskeleton related”, “Translation” and “Mitochondrial proteins”. The confidence score of each interaction is mapped to the edge thickness and opacity. The size of the node relates to the enrichment in log2 fold change (log2FC) over the BioID-GFP background control. The protein abundance in the BioID2-CLUH sample is illustrated by a color scale and corresponds to the specific spectral count normalized to the protein size. Proteins with mitochondrial targeting sequences (MTS) according to Uniprot database are highlighted in red.

To recapitulate, our BioID data on three different cell types strongly suggest that CLUH is transiently interacting or is proximal to nuclear encoded mitochondrial and this vicinity may be related to its function in the cell.

### CLUH promiscuity to CPMPs depends on its TPR domain

To investigate further the link between CLUH and mitochondrial proteins, we engineered *CLUH* knockout mutant cells (*CLUH* KO) using CRISPR/Cas9 technology to delete genomic regions leading to gene inactivation in both HCT116 and mESC backgrounds (Figure S6A and S6B). The generated homozygous HCT116 *CLUH* KO cells showed a complete depletion of the protein (Figure S6C, 5A) and recapitulated previously described clustered mitochondria phenotype (Figure S6G) as well as the proliferation defect (Figure S6H) [11, 12]. Likewise, generated homozygous mESC *CLUH* KO cells (Figure S6D, S6F, S6G) display a clustered mitochondrial phenotype that is more discernable when the cells are differentiated (Figure S6G, S6I), likely due to the immature state of mitochondria in pluripotent mESC cells [19]. The TPR domain of CLUH being important for stable protein-protein interactions, we tested whether it is also required for CLUH promiscuity to CPMPs. To be in condition of absence of endogenous CLUH, we transduced HCT116 *CLUH* KO cells and selected cells stably expressing BioID2-tagged wildtype human CLUH (BioID2-3xHA-CLUH) or CLUH lacking TPR domain (BioID2-3xHA-CLUHΔTPR). We then induced the biotinylation for 24 hours, pulled-down labeled proteins and revealed selected CPMPs by western blot (Figure 5B). All the tested proteins (LRPPRC, IMMT, HSPA9 and ATP5A) were successfully biotinylated by the wildtype human CLUH construct and pulled down on streptavidin beads, recapitulating the MS-analyzed BioID data. Interestingly, the CLUHΔTPR mutant did not biotinylate any of the tested CPMPs, indicating that the TPR domain is required for CLUH proximity to these mitochondrial proteins.

**Figure 5:**
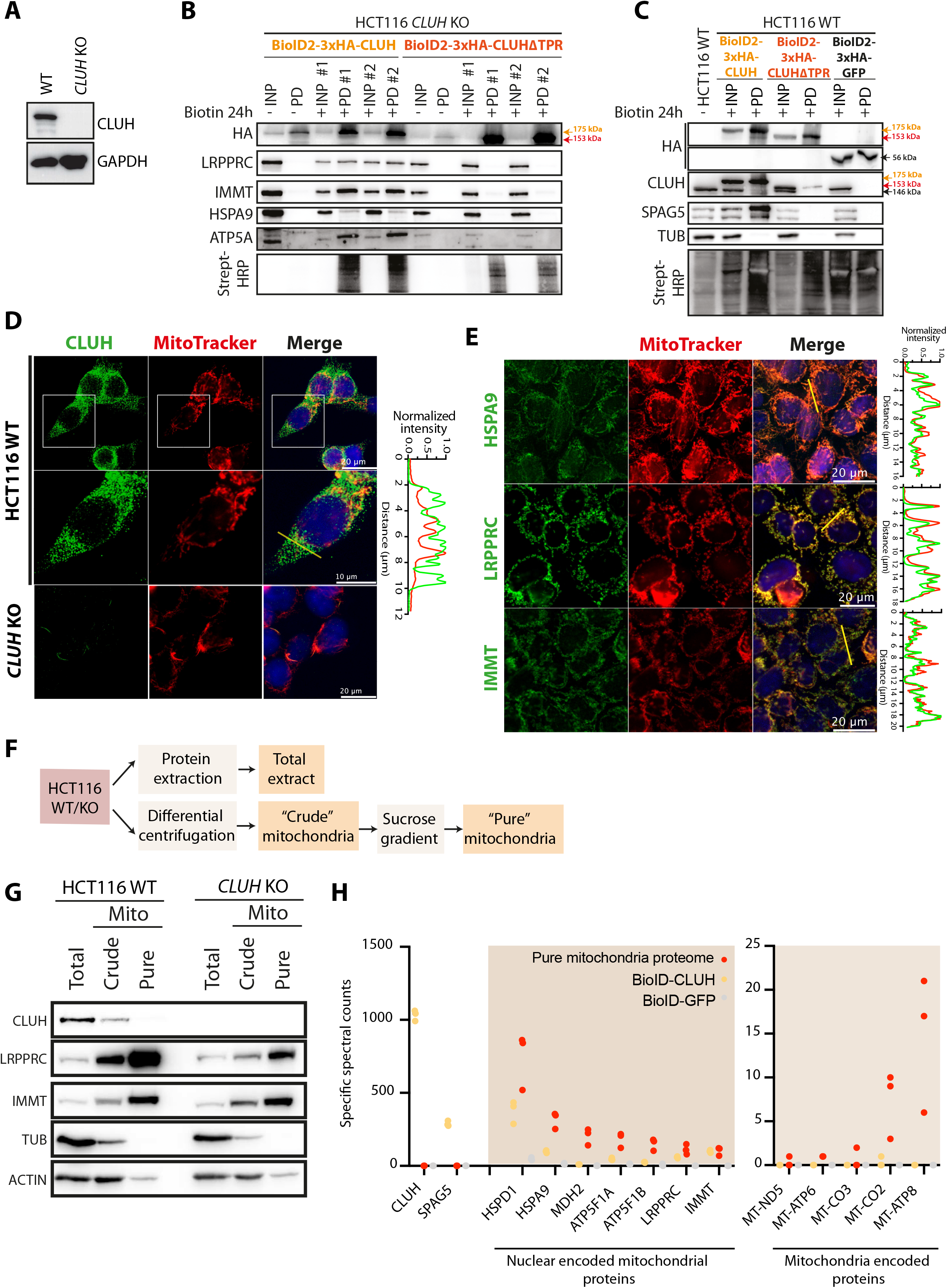
CLUH proximity to mitochondrial proteins occurs in the cytosol and requires the TPR domain. **(A)** Western blot showing the expression of CLUH in HCT116 cells wild type (WT) and knockout for *CLUH* (*CLUH* KO). Indicated proteins are revealed using specific antibodies. **(B-C)** Western blot analysis of BioID experiments performed on *CLUH* KO cells **(B)** and on WT HCT116 cells **(C)** transduced to stably express BioID2-3xHA-CLUH, BioID-3xHA-CLUHΔTPR or BioID2-3xHA-GFP proteins. The proximity labeling is performed for 24 hours in the presence of 50 µM biotin in the culture medium. Biotinylated proteins are pulled down (PD) from total input extracts (INP) using streptavidin-coupled magnetic beads. Replicate experiments are identified with #1 and #2. The constructs are revealed with anti-HA antibody (HA). CPMPs (LRPPRC, IMMT, HSPA9 and ATP5A) and other indicated proteins are revealed using specific antibodies. The size of the endogenous CLUH (black), wildtype transgene (orange) and the delta-TPR mutant (red) is indicated. Biotinylated proteins are revealed using HRP-coupled streptavidin. **(D-E)** Immunofluorescence confocal imaging on HCT116 fixed cells showing the subcellular localization of CLUH **(D)** and three CLUH proximal proteins **(E)**. The endogenous proteins are detected using specific primary antibodies and revealed using Alexa488-coupled secondary antibodies (green). Mitochondria (red) are labeled using MitoTracker™ Red CMXRos. Nuclei (blue) are stained with Hoechst. *CLUH* KO cells are used as control. The fluorescence profile of the red and green channel over the indicated pixel lines are shown on the right. The fluorescence signal is normalized to the highest value for each channel. **(F)** Schematic representation of the experimental workflow to obtain both “crude” and “pure” mitochondria. **(G)** Western blot on total “crude” and “pure” mitochondrial fractions from wild-type and *CLUH* KO HCT116 cells. Indicated proteins are revealed using specific antibodies **(H)** Scatter plot showing the abundance (specific spectral counts) of nuclear and mitochondrial encoded proteins identified by MS from pure mitochondrial fraction (red), BioID2-CLUH (orange) and GFP-BioID2 (grey) samples. Each dot represents a biological replicate samples.

It is intriguing to note that although BioID-3xHA-CLUH co-immunoprecipitated with the endogenous CLUH protein in a TPR dependent manner (Figure 3D), it is no able to biotinylate its endogenous partner when self-interacting (Figure 5C). Indeed, we only detected the self-biotinylated BioID-3xHA-CLUH protein, the endogenous one being presumably not accessible for biotinylation. On the other hand, CLUH-SPAG5 TPR-dependent interaction is revealed using both co-IP and proximity biotinylation approaches (Figures 3D and 5C). This suggests that CLUH self-interaction may not be direct and would involve a heterogenous protein complex comprising more than a single CLUH protein.

### CPMPs interact transiently with CLUH before being imported into mitochondria

CLUH has been described as a cytosolic protein [11]. Nuclear encoded mitochondrial proteins are translated in the cytoplasm and rapidly imported into mitochondria avoiding cytosolic accumulation [32, 33]. To clarify the location of CLUH-induced biotinylation of CPMPs we first verified where the proteins accumulate in our cells. We performed immunofluorescence localization of CLUH (Figure 5D) and of some CPMPs (Figure 5E) in HCT116 wild-type cells. CLUH accumulates in the cytoplasm and do not colocalize with the mitotracker signal, indicating that the protein does not accumulate inside mitochondria. On the other hand, HSPA9, LRPPRC and IMMT clearly accumulates inside mitochondria and show no cytosolic localization. The diverse localization of CLUH and CPMPs was also confirmed by purifying mitochondria on sucrose gradient and analyzing the protein content by western blot (Figure 5F, 5G, S6J). The “crude” fraction, enriched in mitochondrial proteins, is further purified to obtain the “pure” fraction strongly depleted in cytosolic proteins. Despite a weak cytosolic contamination shown by ACTIN (Figure 5G) and GAPDH (Figure S6J) signals, the “pure” mitochondrial fraction is considerably enriched in mitochondrial proteins (IMMT, LRPPRC, GLS) and depleted in CLUH protein compared to the total extract. To be more exhaustive, the pure mitochondrial proteome was analyzed by LC-MS/MS (Table S6) and the abundance of identified proteins compared to the proteins identified in BioID experiment (Figure 5H). Due to the proteome-coverage limitation, only the 1503 most abundant proteins were detected of which 63.2% are annotated in Mitocarta 3.0 database [2] and include 41 of the 43 BioID-identified CPMPs. We also detected 5 out of the 13 proteins encoded by the mitochondrial genome. Importantly, CLUH protein was not detected in the proteome, corroborating the immunofluorescence and western blot observations. We excluded the presence of non-detectable fraction of CLUH inside mitochondria since none of the mitochondria encoded proteins were biotinylated in BioID experiment. Therefore, CLUH proximity to CPMPs, revealed by BioID, occurs most likely in the cytosol before the import inside mitochondria. To rule out the possibility of an accumulation of biotinylated CPMPs in the cytoplasm, we analyzed the protein coverage of the most abundant CPMPs in MS BioID data to detect the presence of an MTS (Figure S6K). None of the analyzed mitochondrial proteins included the MTS-containing N-terminal region, indicating that the proteins are first biotinylated, imported, matured and then accumulate inside mitochondria.

### CLUH proximity-interactors accumulate over time

To verify if the identified CLUH-CPMPs cytosolic transient-interactions are related to a dynamic biological process, we performed a time-course proximity labeling experiment in order to capture the variation of CLUH proximity-interactome over time. We took advantage of the increased biotinylation efficiency of the TurboID enzyme [25, 34] to label CLUH proximal proteins for 30 minutes and 16 hours (Figure 6A). We generated HCT116 stable cell lines expressing human CLUH or GFP proteins fused to TurboID in N-terminal (Figure S7A). As for the BioID, the labeling was initiated by the addition of biotin in the culture medium and the cells collected at the two time points displaying increasing global biotinylation levels (Figure S7B). Biotinlylated proteins are then enriched on streptavidin-coated beads and analyzed by LC-MS/MS (Table S7). We identified 106 and 244 significantly enriched proteins compared to the GFP control at respectively 30 mins and 16 hours of labeling (Figure S7C, S7D). Alike the BioID experiment, the ontology analysis revealed an enrichment in terms associated to mitochondria and to respiration (Figure S7E). Although the same functional categories came out at both time points, the enrichment is more prominent at 16 hours.

**Figure 6:**
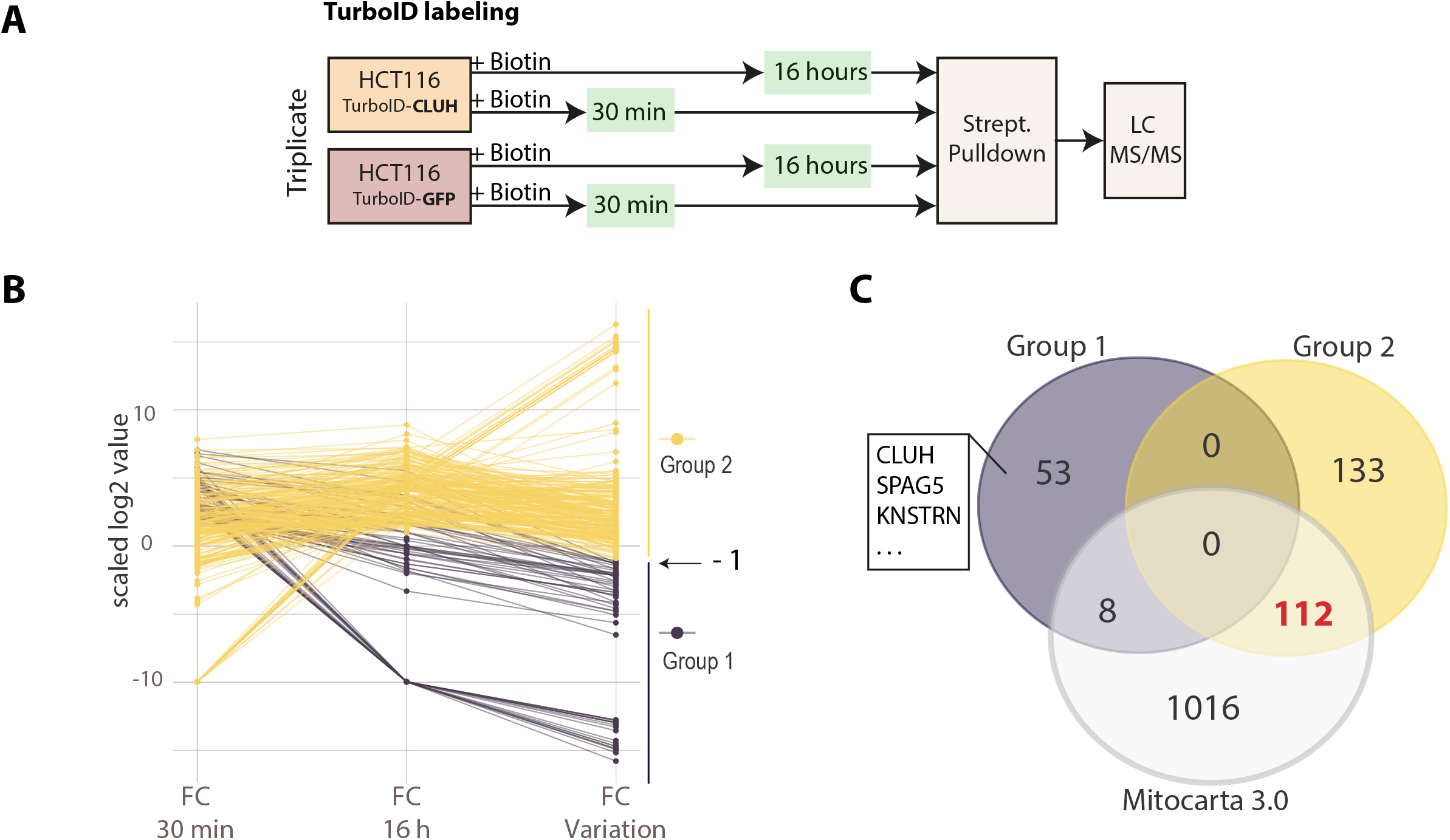
CLUH-proximal mitochondrial proteins accumulate overtime. **(A)** Schematic representation of the TurboID experimental design using HCT116 cells stably expressing the TurboID protein fused to CLUH or GFP proteins. The proximity labeling is performed for 30 minutes or 16 hours in the presence of 50 µM biotin in the culture medium. Biotinylated proteins, from both the specific (TurboID-CLUH) and control (TurboID-GFP-BioID2) samples, are isolated using streptavidin-coupled magnetic beads and identified by LC MS/MS. **(B)** Parallel coordinates plot comparing the fold change (FC) enrichment of TurboID identified proteins at 30 mins and 16 hours (n=3 and threshold: FDR<0.1 and FC >2, Figure S5 C and D). The FC variation corresponds to the ratio of FC between 30 min and 16h. The y-axis corresponds to the log2 transformed values. Group 1 (purple) is define as proteins with a FC ratio < 0.5 and group 2 (yellow) as protein with FC variation >=0.5. **(C)** Venn diagram showing the intersection between group 1, group 2 and human mitochondrial proteins listed in Mitocarta 3.0 database.

We compared the fold change (FC) enrichment over GFP control of all significantly enriched proteins at both time points in order to capture a variation pattern (Figure 6B). We empirically defined two groups of proteins delimited by a FC variation of 0.5 between 30 minutes and 16 hours. Group 1 contains proteins showing a reduced enrichment over time while group 2 contains proteins showing a relatively stable or increased enrichment over time. Interestingly, group 2 contained 93,3% of identified mitochondrial proteins while the group 1 contained only 6,6% mitochondrial proteins, the bait CLUH and previously co-IP identified stable interactants such as SPAG5 and KNSTRN (Figure 6C, S7F). This distribution could be explained by the fact that stable interactants are labelled once and do no change much over the time frame of the experiment while mitochondrial proteins are continuously translated, biotinylated by CLUH, and rapidly imported into mitochondria. The transient contact with CLUH and the accumulation in a separated compartment favor interactions with newly synthetized proteins.

Altogether our data indicate that CLUH transiently interacts with nuclear encoded mitochondrial proteins prior their import and accumulation inside mitochondria.

This transient interaction is presumably related to a dynamic biological process that may be related to their translation or import.

### CLUH is associated to mRNAs coding for CPMPs through its TPR domain with no impact on their translation efficiency

CLUH has been previously described as an RNA binding protein. Interestingly, all the mRNAs coding for CPMPs have been previously identified as being bound by CLUH in HeLa cells [11]. We verified CLUH association to the mRNAs of 10 randomly selected CPMPs by performing a UV crosslinked RNA-immunoprecipitation (RIP) experiment on endogenous CLUH protein in wild type HCT116 (Figure 7A). The CLUH enriched mRNAs were quantified by RT-qPCR and compared to a *CLUH* KO background control sample. Whilst there was no enrichment for EIF5A and SUB1 mRNAs coding for cytosolic proteins, we observed a clear enrichment for the 10 mRNAs coding for CPMPs, confirming the specific CLUH binding to these mRNAs. Since CLUH proximity to CPMPs is dependent on its TPR domain, we wanted to verify if this domain was also required for its association to cognate mRNAs. We used the previously generated knockout-rescued stable cell lines (Figure 5B) to perform, as previously, a RIP-qPCR experiment using antibodies directed against CLUH protein (Figure 7B). We successfully pulled down mRNAs coding for CPMPs in *CLUH* KO cells rescued with the wild-type CLUH protein, compared to cells expressing the GFP control protein. However, we observed no specific enrichment in cells expressing the CLUHΔTPR construct, indicating that the TPR domain is required for CLUH binding to mRNAs. Overall, the TPR domain seems to be important for CLUH interaction with both CPMPs and their mRNAs. Such a proximity may be happening during the translation of CPMPs. Therefore, we verified the impact of CLUH deletion on the translation efficiency of CPMPs, by quantifying the mRNAs in the polysomal fraction of both wild-type and *CLUH* KO HCT116 cells (Figure 7C, 7D). We first isolated the polysomes from both cell lines on a sucrose gradient (Figure 7C) and isolated associated RNAs (Figure S8A). We then quantified and analyzed the enrichment over the input of mRNAs coding for the 10 previously analyzed CPMPs, for two cytosolic proteins (*GAPDH* and *SUB1*) and for one mitochondria-encoded protein (*MTCO2*). Surprisingly, we observed no difference in mRNA enrichment between wild-type cells and cells lacking CLUH, meaning that the absence of CLUH has no impact on their translation efficiency (Figure 7D). Evidently, there was no enrichment for *MTCO2* as we isolated principally cytosolic polysomes. The impact of CLUH on CPMPs steady-state levels in HCT116 was also analyzed by comparing the proteome of isolated “pure” mitochondria (Figure 5G) from wild-type and *CLUH* KO cells (Figure 7E, Table S6). We observed no obvious bias in the abundance of mitochondrial proteins in the absence of CLUH, using z-scoring method from three replicate samples (Figure 7E). This observation was also confirmed by analyzing individual proteins in wild-type and *CLUH* KO HCT116 cells by western blot (Figure S8B). Interestingly, the analysis of the proteins extracted from the polysomal fractions by western blot, revealed the presence of CLUH in fractions containing translating ribosomes (Figure 7F). This strongly suggest that CLUH, as an RNA binding protein, remains associated with the mRNAs coding for CPMPs during their translation. This association to mRNAs in translation may explain its proximity to newly synthetized mitochondrial proteins as observed in the BioID experiment.

**Figure 7:**
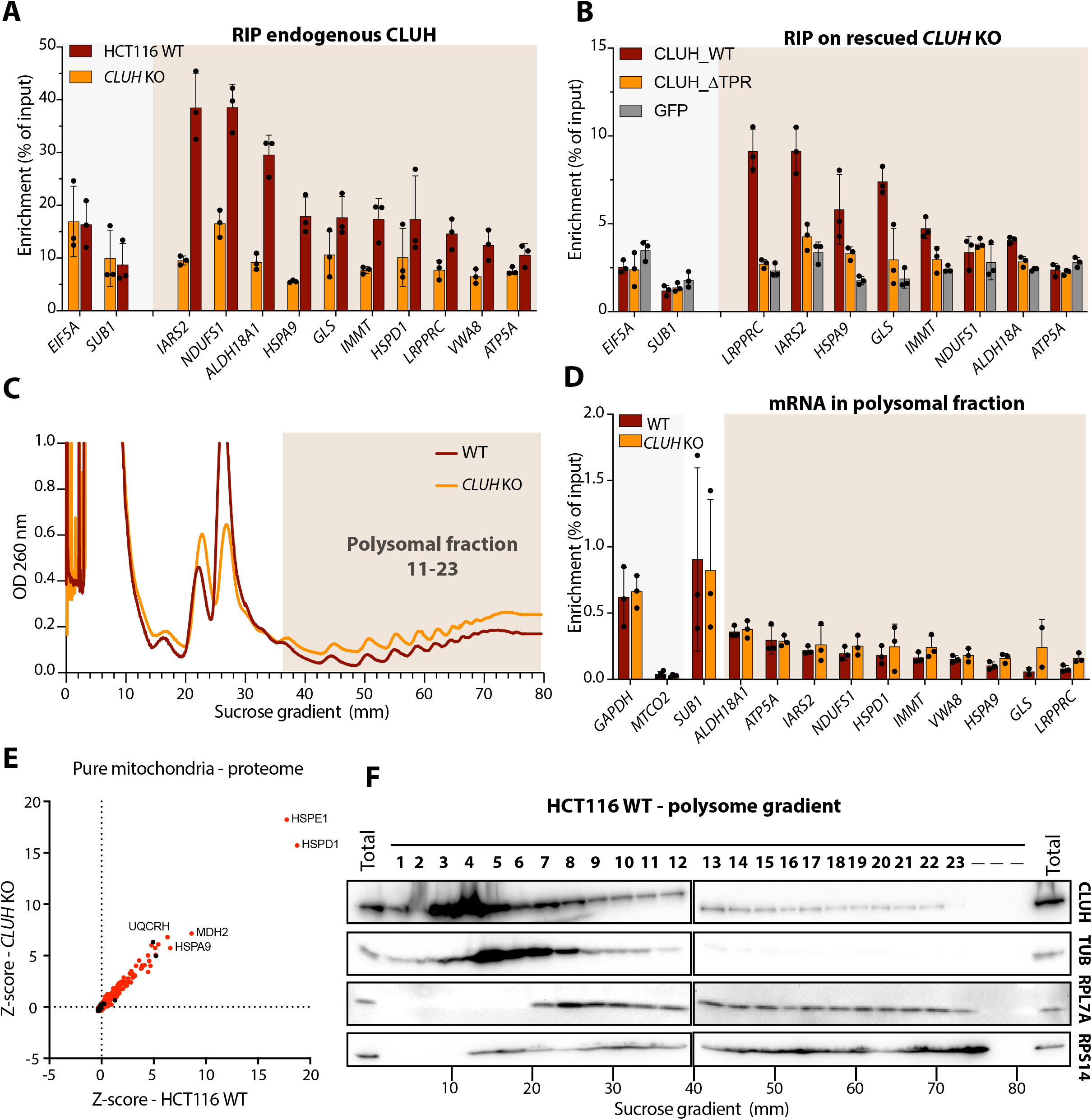
CLUH binds mRNAs coding for mitochondrial proteins and do not affect their translation efficiency. **(A)** RT-qPCR analysis of RNA immunoprecipitation (RIP) experiments performed on the endogenous CLUH protein in wild-type HCT116 (red) and *CLUH* KO (orange) cells. **(B)** RT-qPCR analysis of RIP experiments performed on rescued HCT116 *CLUH* KO cells. The cells are transduced to stably express either the tagged wildtype CLUH (CLUH_WT, red) or the tagged mutant CLUH (CLUH_ΔTPR, yellow). Cells expressing a tagged GFP protein (GFP, grey) are used as background control. **(A-B)** The CLUH associated mRNAs are enriched using CLUH-specific antibodies and measured by RT-qPCR. The enrichment of specific mRNA (normalized to *GAPDH* levels) is calculated relative the input sample (% of input). mRNAs coding for CPMPs are highlighted in orange. The error bar corresponds to the standard deviation of three independent experiments. The average value for each replicate is indicated by a dot. **(C)** Representative graphs of polysome profilings of WT and *CLUH* KO HCT116 cells. The y-axis corresponds to the absorbance at 260 nm and the x-axis to the distance in the sucrose gradient. The polysomal fractions used for further experiments are highlighted in orange. **(D)** RT-qPCR analysis of mRNA in polysomal fractions from the WT and *CLUH* KO cells. The enrichment of specific mRNA (normalized to *EIF5A* levels) is calculated relative the input sample (% of input). The error bar corresponds to the standard deviation of three independent experiments. The average value for each replicate is indicated by a dot. mRNAs coding for CPMPs are highlighted in orange. **(E)** Scatter plot comparing the abundance of all proteins identified by mass spectrometry in pure mitochondrial fraction (see Figure 4F) of *CLUH* KO and WT HCT1116 cells. The x-axis and the y-axis show to the Z-score of the mean abundance of each protein in the WT HCT116 and *CLUH* KO samples, respectively. The abundance of each protein is calculated by dividing the mean spectral count of three replicate samples by the protein size. Proteins found in the Mitocarta 3.0 database are shown in red. **(F)** Representative western blot analysis of polysome profiling of WT HCT116 cells. Each fraction corresponds to a about 3.3 mm of the sucrose gradient and are numbered from 1 to 23. Total protein extracts are used as controls. Indicated proteins are revealed using specific antibodies.

Although CLUH has been previously reported to affect the decay of some targeted mRNAs [12], we did not observe any impact on the stability of several mRNAs coding for CPMPs in HCT116 cells (Figure S8C).

### The subcellular localization of mRNAs coding CPMPs is altered in the absence of CLUH

The current view on the translation of nuclear encoded mitochondrial proteins points toward a cytosolic translation and a translation located at the mitochondrial surface [3, 4]. In mammals, this model is supported by the recent discovery of mRNAs located at the mitochondrial surface in a translation dependent manner [6]. Interestingly, almost all the mRNA coding for CPMPs have been described as being specifically enriched at the mitochondrial outer membrane (OMM) by APEX-seq (Figure 8A). To verify whether CLUH may be involved in mRNA localization at the OMM, we isolated “crude” mitochondria fraction from both HCT116 wild-type and *CLUH* KO cells (Figure 5G) and extracted the associated RNAs. The “crude” fraction was preferred over the “pure” in order to preserve the mRNA association to the OMM. We quantified and compared mRNA enrichment in the mitochondrial fraction over the input in wild-type and *CLUH* KO cells (Figure 8B). We observed a strong enrichment of mitochondria encoded *MTCO2* in both samples and weak enrichment for *SUB1* and *GAPDH* mRNAs (coding for cytosolic proteins), confirming the mitochondrial enrichment and the cytosolic depletion. Surprisingly, while the enrichment of half of the mRNAs coding for CPMPs did not change, we observed a higher enrichment of *GLS*, *HSPA9*, *LRPPRC*, *HSPD1* and *IMMT* mRNAs in the *CLUH* KO mitochondrial fraction compared to the wild-type sample. CLUH being associated to CPMPs in translation, we attempted to isolate polysome from the “crude” mitochondria enriched fraction (Figure 8C, 5G) and verify their translation efficiency. Despite the limiting material we extracted both proteins and RNAs from the polysomal fractions and confirmed the presence of ribosomal subunits (Figure S8A, S8B). As for the total cytosolic samples, we could detect CLUH proteins in the polysomal fractions from the “crude” mitochondria enriched samples. We then analyzed by RT-qPCR the mRNA enrichment in the isolated polysomes. Interestingly, all the mRNA presenting a higher enrichment in the mitochondrial fraction (Figure 8B) in the absence of CLUH showed also a higher enrichment in the polysomal fraction, suggesting a higher translation efficiency (Figure 8D). The absence of CLUH affected the translation efficiency of some mRNAs in mitochondrial polysomes but not in total polysomes. Therefore, in terms of proportion, CLUH-affected mRNAs coding for CPMPs at the mitochondrial surface are negligeable compared to those translated in the cytosol (Figure 7D). Taken together our data indicate that CLUH may be involved in the subcellular localization of a small fraction of specific mRNAs near mitochondria and may affect their localized-translation efficiency.

**Figure 8:**
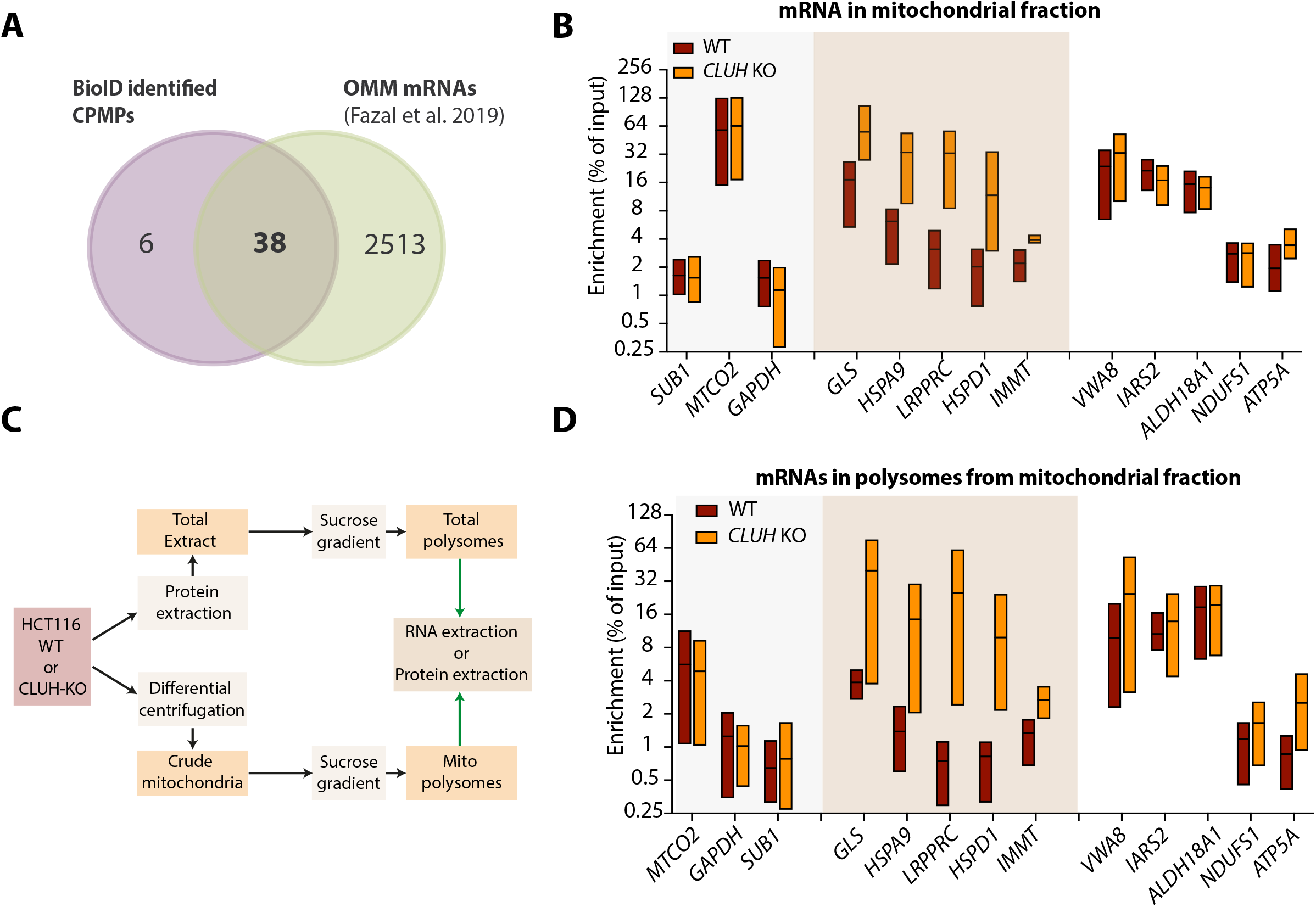
CLUH affects the subcellular distribution of some mRNAs coding for mitochondrial proteins. **(A)** Venn diagram depicting the intersection between mitochondrial CPMPs (see Figure 4E) and the mRNAs identified by APEX-seq at the mitochondrial outer membrane (OMM) [6]. Mitochondrial proteins are selected according to their presence in the Mitocarta 3.0 database. The OMM localized mRNAs correspond to OMM biotinylated mRNA in the presence of cycloheximide significantly enriched above the background (FDR < 10%) and positively enriched (log2FC >0) over non-specific cytosolic biotinylation. **(B)** Graph showing RT-qPCR analysis of mRNA enrichment in crude mitochondrial fraction from *CLUH* KO (orange) and WT HCT116 cells (red). The enrichment of specific mRNA (normalized to *EIF5A* levels) is calculated relative to the total RNA from non-fractionated input sample (% of input) and shown on log2 scaled y-axis. The box summarizes triplicate experiments, showing the upper, the lowest and mean enrichment values. mRNAs showing the highest variation between WT and *CLUH* KO are highlighted in orange. **(C)** Schematic representation of the experimental workflow to extract RNA and proteins from polysomal fractions from both total and “crude” mitochondrial fractions. **(E)** Graph showing RT-qPCR analysis of mRNA enrichment in the polysomes from crude mitochondrial fraction in *CLUH* KO (orange) and WT HCT116 cells (red). The enrichment of specific mRNA (normalized to *EIF5A* levels) is calculated relative the total RNA from non-fractionated input sample (% of input) and shown on log2 scaled y-axis. The box summarizes triplicate experiments, showing the upper, the lowest and mean enrichment values. mRNAs showing the highest variation between WT and *CLUH* KO are highlighted in orange.

## MATERIAL AND METHODS

### Cell lines and cell culture

HCT116 line (ATCC^®^ CCL-247^™^) and derived stable cell lines were grown in standard DMEM (D6429, Sigma-Aldrich) with 10% fetal bovine serum (FBS, Gibco) and 1% Pen-Strep (Sigma-Aldrich). E14 mESC line (ATCC^®^ CRL-1821^™^) and derived lines were grown in DMEM (D6429, Sigma-Aldrich), containing 15% of FBS (Gibco), 100 U/mL LIF (Millipore), 0.1 mM 2-ß-mercaptoethanol (Gibco) and 1% Pen-strep (Sigma-Aldrich), on 0.2% gelatin-coated plates. All cells were grown at 37°C in 5% CO2 humidified atmosphere. When required, the culture medium was supplemented with 2 µg/mL of puromycin (InvivoGen) and 10 µg/mL of blasticidin (InvivoGen).

### Embryoid body (EB) differentiation

E14 mESC and derived lines were differentiated as previously described [35, 36]. Briefly, the EBs were obtained by culturing the cells in suspension for 6 days in non-tissue culture treated dishes to prevent attachment. After 6 days the EBs were plated on 0.2 % gelatin-coated culture-treated dishes for 4 additional days.

### Gibson Assembly and sgRNA cloning

All the constructs (Table S8) were generated using Gibson assembly method [37]. Primers to amplify fragments for the assembly were designed using NEBuilder Assembly Tool (New England Biolabs). Sequences of the assembled plasmids are available upon request. The sgRNAs for CRISPR/Cas9 genome editing were cloned by annealing and ligating oligonucleotides (Table S9) into BBSI-digested pX459 vector.

### Transfections and transductions

All plasmids transfections were performed in 6-well plates on 70% confluent cells, using PEI method. Briefly, 3 µg of DNA diluted in 100 µL of NaCl 150 mM was mixed with 6 µL of PEI Max 40000 (Polysciences) at 2 mg/mL diluted in 100 µL of NaCl 150 mM. After 30 min incubation at RT, the mix was added into the cells (2 mL medium volume) and the medium changed after 6 hours. SiRNAs were transfected at 60nM final concentration using Lipofectamine 2000 (ThermoFisher scientific) according to manufacturer instructions. The transfection was repeated after 48 hours to increase knockdown efficiency. Used siRNA duplexes are listed in Table S8.

All stable cell lines were produced by transducing the parental line with lentiviral particles followed by antibiotic selection. Lentiviral particles were produced by transfecting 293T cells in 6-well plates with 1.5 µg of vector and 1.5 µg of the packaging plasmids psPAX2 and pVSV-G (in proportions 4:1). Retroviral particles were produced using 1.4 µg of viral vector, 0.250 µg of pAdventage and 1.4µg of packaging plasmids pGAG_pol and pCMV-VSV-G in proportion 1.75:1. The viral supernatant was collected after 48 hours and filtered using a 0.45µM PES filter. Cells were transduced in 6-well plates using 500 µL of viral supernatant and 500µL of fresh medium supplemented with 8µg/mL of polybrene (Sigma-Aldrich). The medium was changed after 24 hours and the selection with the appropriate antibiotic initiated after 48 hours. Cells were maintained under selection for a minimum of 10 days before experimentation.

### CRISPR/Cas9 mediated genome editing

All the knockout lines were generated using paired CRISPR/Cas9 strategy to induce a genomic deletion. The cells were transiently transfected with pX459 vector coding for specific sgRNAs. Transfected cells were selected with puromycin for 2 days at 2µg/mL and monoclonal cell populations were isolated by limiting dilution method. Homozygous knockouts were screened by genotyping PCR, validated by sequencing and western blot. mESCs *Cluh* KO1 (clone C4) and KO2 (clone B12) were generated with sgRNA1/2 and sgRNA3/4 respectively. HCT116 *CLUH* KO cells were generated using sgRNA5/6. The *Cluh* knock-in mESCs 3xHA-CLUH (Clones G6 and G12) and BioID2-CLUH (clones E2 and C2), were generated by transfecting the sgRNA7 together with the adequate homologous recombination template (pNG12 or pNG13). The sequences of all sgRNA and PCR primers are listed in Table S9.

### Co-immunoprecipitation

The co-immunoprecipitations were performed using µMACS HA or GFP Isolation Kit (Miltenyi Biotec) according to manufacturer protocol using 10×10^6^ cells and 50 µL of antibody-coupled magnetic beads in a volume of 1 mL. Both the lysis and washes were done using the supplied Lysis Buffer supplemented with protease inhibitors (cOmplete™ Roche). When required, before adding the beads, the total protein extract was supplemented with 20 µg/mL RNAse A and 50 U/mL of RNAse T1 (RNAse A/T1 Mix, Thermofisher Scientific) and incubated 10 min at 37°C.

### Western blots

The protein extracts were prepared from cell pellets using RIPA lysis buffer (50 mM Tris HCL, pH 7.4; 150 mM NaCl; 0.1% SDS; 0.5% Sodium deoxycholate; 1% triton 100X) supplemented with protease inhibitors (cOmplete™ Roche) 10 mins on ice. The extracts were sonicated 3 times 30 seconds at 20% amplitude and the debris pelleted by centrifugation 10 minutes at 12 000 g. The extracts were quantified using BCA assay (Pierce, Thermofisher Scientific). Proteins were separated on SDS-PAGE and transferred on PVDF or Nitrocellulose membrane and revealed using primary antibodies listed in Table S10. To reveal biotinylated proteins, the membrane was blocked with BSA blocking buffer (PBS, 1% BSA, 0.2% Triton X-100) and incubated 1 hour at RT or 16 hours at 4°C with HRP-coupled streptavidin (RPN1231VS, GE Healthcare) diluted 1:20000 in BSA blocking buffer. The membrane was washed with PBS and ABS blocking Buffer (PBS, 10% fetal bovine serum, 1% triton X-100) for 5 min, before revealing with ECL reagent.

### Split-GFP Interaction assay and Confocal microscopy imaging

For the Split-GFP assay, wild-type HCT116 were transduced with pNG43 expressing SPAG5 fused to the sfGFP1-10 fragment together with pNG45 or pNG47 expressing respectively CLUH and GAPDH fused to mCherry-sfGFP11 fragment. The cells were selected using both 10µg/mL of blasticidin and 2 µg/mL of puromycin in order to select cells expressing two constructs. The localization and the expression of transgenes was analyzed by directly imaging the fluorescence of GFP-and BFP-fusion proteins or by immunodetection using antibodies specific to the V5 tag. HCT116 wt cells were used to analyze endogenous protein localization by immunofluorescence. Cells were grown on 8-well Culture Slide (Falcon) in complete DMEM culture medium or in HBSS (Gibco), if required supplemented with CHX (0.1 mg/mL) or Sodium Arsenite (0.5 mM). Mitochondrial were labeled by growing the cells in presence of 200 nM of MitoTracker™ Red CMXRos (Thermofisher Scientific) for 1 hour. Cells were washed 2 times with fresh medium for 5 minutes and fixed for 15 minutes at RT with PFA 4% in PBS. If immunolabelling was required, the cells were permeabilized for 1 hour using Blocking/permeabilization buffer (1% BSA; 5% Goat serum and 0,25% triton X-100 in PBS 1X) and incubated overnight with primary antibody (Listed in supplementary table S10) diluted in Blocking/permeabilization buffer. After 3 washed with PBS for 5 minutes, the cells were incubated with Alexa Fluor® 488 or Cy5-conjugated secondary antibody (Invitrogen) at 1 : 500 dillution in Blocking/permeabilization buffer for 1 hour at RT and washed again 3 times with PBS for 5 minutes. To label nuclei, the cells are incubated 5 minutes with 1µg/mL of Hoechst 33342 (Thermofisher Scientific) in PBS. The slides were mounted using Shandon™ Immu-Mount™ medium. Samples expressing eGFP-, mCherry-or BFP-fusion proteins as well as immuno-detected proteins were imaged on Zeiss LSM 780 confocal microscope using ZEN software.

The images were analyzed using Fiji software [38] and Aggrecount plugin [39]to quantify Split-GFP signal aggregates. The microscopy figures were mounted using FigureJ [40] plugin.

### BioID proximity labeling

The BioID experiment was based on the previously described protocol by [24] with small changes, using HCT116 cells stably expressing BioID2-mCLUH and genome edited mESCs expressing endogenously tagged BioID2-CLUH. Cells stably expressing GFP-BioID2 fusion are used as non-specific control. The biotin labeling was performed for 24 hours in presence of 50 µM of biotin (Sigma Aldrich) in the complete culture medium. About 40 million of cells (4x 10-cm plates), per experiment, were washed in PBS and lysed in 2,4 mL of lysis buffer (50 mM Tris HCL, pH 7.4; 500 mM NaCl; 0.4% SDS; EDTA 5mM;1 mM dithiothreitol) and sonicated 3 times at 20 % amplitude for 20 seconds. Triton X100 was added to a final concentration of 2% and the sample diluted 2 times with 50 mM Tris HCL, pH 7.3. The sample was centrifuged for 10 minutes at 12 000 g and the supernatant kept to pulldown biotinylated proteins by adding 200 µL of streptavidin-coupled magnetic beads slurry (Streptavidin Mag Sepharose, GE Healthcare) previously washed and equilibrated. Samples were put on a rotating wheel overnight at 4°C. The beads were washed twice for 5 min at RT on rotating wheel with wash buffer 1 (2% SDS), once with wash buffer 2 (0.1% deoxycholate; 1% Triton X-100; 500 mM NaCl; 1 mM EDTA; and 50 mM HEPES; pH 7.5), once with wash buffer 3 (250 mM LiCl; 0.5% NP-40; 0.5% deoxycholate; 1 mM EDTA; and 10 mM Tris, pH 8) and once with wash buffer 4 (50 mM Tris, pH 7.4). All buffers are supplemented with protease inhibitors (cOmplete™ Roche). Beads were washed again 2 times with 50 mM NH4HCO3. A fraction of the beads (5%) was boiled in Laemmli buffer for western blotting and the remaining beads were analyzed by LC-MS/MS.

### TurboID proximity labeling

TurboID proximity labeling is based on previously described protocol [34]. Briefly, HCT116 stable cell line expressing TurboID-CLUH and TurboID-GFP were grown for 30 minutes and 16 hours in presence of 50 µM of biotin (Sigma Aldrich). About 20 million cells (two 10-cm plates) were first washed two times with ice-cold PBS and lysed in 1 mL of RIPA buffer (50 mM Tris HCL, pH 7.4; 150 mM NaCl, 0.1% SDS, 0.5% Sodium Deoxycholate, 1% triton 100X) for 10 minutes on ice. The extract is sonicated 3 times at 20 % amplitude for 20 seconds and cleared by centrifugation at 12 000 g for 10 minutes. The supernatant was kept to pulldown biotinylated proteins by the addition of 100 µL of streptavidin-coupled magnetic beads slurry (Streptavidin Mag Sepharose, GE Healthcare) previously washed and equilibrated. Samples were put on rotating wheel overnight at 4°C. Beads were washed 5 minutes at RT on rotating wheel, twice with RIPA buffer, once with 1M KCL, with 0.1M Na2CO3 and with 2M urea in 10mM Tris-HCL pH8. Beads are washed again twice with RIPA buffer and with 50 mM NH4HCO3. A fraction of the beads (5%) was boiled in Laemmli buffer for western blotting and the remaining beads were analyzed by LC-MS/MS.

### LC-MS/MS

For mass spectrometry analyses, proteins eluted in Laemmli buffer from immunoprecipitations and protein extracts from mitochondrial proteomes were prepared as previously described [41]. Briefly, eluted immunoprecipitated proteins or 5 micrograms of mitochondrial proteomes were precipitated with two cycles of 0.1 M ammonium acetate in 100% methanol overnight precipitations, reduced with 5mM dithiothreitol (10 min, 95°C) and alkylated with 10mM iodoacetamide (30 min, RT, in the dark). After quenching with 5 mM dithiothreitol, proteins were digested overnight with sequencing-grade porcine trypsin (Promega, Fitchburg, MA, USA).

For the proximity labeling experiments, magnetic beads were extensively washed in 50 mM ammonium bicarbonate and proteins were digested directly on the beads in 2 consecutive steps with sequencing grade trypsin.

Peptides generated after trypsin digestion were analyzed by nanoLC-MS/MS on a QExactive + mass spectrometer coupled to an EASY-nanoLC-1000 (Thermo-Fisher Scientific, USA). Peptides were identified with Mascot algorithm (Matrix Science, London, UK): the data were searched against the Swissprot updated databases with Mus musculus or Homo sapiens taxonomies using the software’s decoy strategy. Mascot and Swissprot versions used for each experiment are mentioned in results tables. Mascot identifications were imported into Proline 1.4 software [42] where they were validated using the following settings: PSM score <=25, Mascot pretty rank < = 1, FDR < = 1% for PSM scores, FDR < = 1% and for protein set scores. The total number of MS/MS fragmentation spectra was used to quantify each protein. Mass spectrometry proteomics data have been deposited to the ProteomeXchange Consortium via the PRIDE partner repository with identifiers PXD 027158 and PXD027122.

### Analysis of proteomic data

The enrichment analysis of co-IP, BioID and TurboID experiments was performed using SAINTxpress software [26] on label-free spectral count data. Three replicate experiments were analyzed per sample and a confidence score was assigned to the enrichment of proteins for specific Bait samples over non-specific background samples. All proteins passing the selected of Log2 fold change (Log2FC>2) and false discovery rate (FDR<0.1) thresholds were considered as significantly enriched. The functional enrichment analysis was performed using the R package gprofiler2 [43]. Gene network analysis and representation of BioID identified protein was done using the Cytoscape StringApp [44].

### Mitochondria purification

Mitochondria-enriched “crude” extracts and “pure” mitochondria were prepared as described previously [45]. Briefly, cells form ten 10-cm plates (about 10^7^ cells) at 80% confluency are lysed mechanically with sonication (3 times 10 seconds at 30%, BioBlock vibracell 75115) in 2 mL of MTE buffer (270mM D-mannitol, 10mM Tris base, 0.1 mM EDTA, pH adjusted to 7.4) supplemented with protease (cOmplete™ Roche) and RNAse (RNaseOut, Thermofisher Scientific) inhibitors. After a low-speed centrifugation at 700 g for 10 minutes at 4°C to remove debris, the supernatant is centrifuged again for 10 minutes at 15 000 g and 4°C to pellet “crude” mitochondria. The obtained mitochondria-enriched fraction is used for RNA extraction, protein extraction or polysome fractionation. To obtain “pure” mitochondria, the pellet is delicately washed, resuspended in 800µL of MTE buffer and loaded on a discontinuous sucrose gradient (1mL of 1.7 M and 1.6 mL of 1.0 M sucrose) followed by an additional 800µL of MTE. The centrifugation is performed on a SW60Ti rotor at 40 000 g for 22 minutes at 4°C (Beckman Optima L-90K Ultracentrifuge). The mitochondrial fraction (400µL) is collected at the interphase of sucrose layers, washed with 1.1 mL of MTE buffer and centrifuged 10 minutes at 15 000 g and 4°C to pellet “pure” mitochondria. The sample is resuspended in RIPA buffer for protein analysis by western blot or LC-MS/MS.

### Polysome profiling analysis

The polysome fractionation was performed according to a previously published protocol [46]. Briefly, cellular pellet from a 10-cm plate or enriched “crude” mitochondria pellet obtained from ten 10-cm plates are used to isolate polysomes. All cells are collected at 80% confluency and 100µg/mL cycloheximide is maintained in all washes and buffers. The pellet is resuspended in 500µL of Lysis buffer (50mM KCL; 20mM Tris HCL, pH 7.4; 10 mM MgCl2; 1% Triton X100; 1mM 1,4-dithiothreitol; 0.5% sodium deoxycholate; 100µg/mL cycloheximide) supplemented with protease (cOmplete™ Roche) and RNAse (RNaseOut, Thermofisher Scientific) inhibitors, incubated 15 min on ice and centrifuged for 5 minutes at 2 000 g and 4°C. The supernatant is again centrifuged for 5 minutes at 13 000 g and 4°C. The sample is loaded on a 7-47% sucrose gradient and centrifuged at 260 808 g on a SW41Ti rotor for 90 minutes. Fractions collected over the whole gradient and analyzed at 260/280 nm using Gradient Fractionator (Biocomp). Both RNA and proteins are extracted from each fraction using 500µL of Tri reagent (MRC).

### RNA extraction and quantification by RT-qPCR

Total cellular RNA was extracted from a pellet of 1-10 million cells using Tri Reagent® (MRC) according to manufacturer’s protocol. The purified RNA was quantified using Nanodrop2000 (Thermofisher Scientific) and its quality verified on agarose gel. RNA from RIP samples, from mitochondrial fractions and from polysomal fractions was extracted using 500µL of Tri Reagent following manufacturer protocol. The reverse transcription was performed in a final volume of 20 µL using Superscript IV (Thermofisher scientific) and Random hexamers on either 1-2 µg of total RNA, 60 ng of RNA from mitochondrial/polysomal fractions, or on using 11µL of RIP elution and input samples. The quantitative PCR was performed on a Light Cycler 480 (Roche) using 2 μl of the diluted cDNAs (1:5) and Takyon Blue Master Mix (Eurogentec). All primers used are listed in Table S9. The relative quantities and differences between samples are calculated using the 2^−ΔΔCT^ method using *GAPDH* mRNA levels as normalizer.

### mRNA stability assay

Wild type and *CLUH* KO HCT116 cells were plated in 6-well plates and were treated with 10 µg/mL of actinomycin D for 0, 4, 8, 16 and 24 hours. The samples were collected at the different time points by removing the medium and by resuspending the cells in 1 mL of Tri reagent (MRC). The RNA is extracted and quantified by RT-qPCR. The RNA decay over time is calculated relative to *GAPDH* mRNA levels and the t=0 time point.

### RNA immunoprecipitation

Cells are grown on 10 cm plates until 80-90% confluency and UV-crosslinked at 400 mj/cm^2^ (Hoefer, UV-crosslinker). After a wash with PBS the cells are lysed with 1 mL of cold lysis buffer (50mM Tris-HCl pH7.4, 100mM NaCl, 1mM MgCl2, 0.1 mM Ca Cl2, 1% IGEPAL CA-630, 0.1% SDS and 0.5% sodium deoxycholate) supplemented with 8 U of Turbo DNAse I, protease inhibitor (cOmplete™ Roche) and RNAse inhibitor (RNaseOut, Thermofisher Scientific). The lysate is incubated 15 minutes on ice and cleared by centrifugation 10 minutes at 11 000 g. 1% of the input sample is taken into 140 µL of RIP elution buffer (10 mM EDTA, 100 mM Tris– HCl (pH 8.0), 1% sodium dodecyl sulfate (w/v)) supplemented with RNAse inhibitor (RNaseOut, Thermofisher Scientific). The remaining sample is incubated with 0,6 µg of anti-CLUH antibody (NB100-93306, Novus biologicals) 2 hours on rotating wheel at 4°C. Complexes are immunoprecipitated with 25µL of tRNA/BSA blocked protein A-coupled dynabeads (Thermofisher Scientific) 2 hours at 4°C. Beads are washed once with 1 mL lysis buffer 5 minutes at 4°C and once with 1 mL of HighSalt buffer (50µL Tris-HCl pH7.4, 1M NaCl, 1% IGEPAL CA630, 0.1% SDS, 0.5% Sodium deoxycholate and 1mM EDTA) containing RNAse and protease inhibitors. Beads are resuspended in 75µL of RIP elution buffer with 40U of RNAse inhibitors and incubated 10 minutes at 37°C. The supernatant is retrieved, and elution repeated once and pooled. Both RIP and Input samples are supplemented with 6µL of NaCL 5M and 20µg of proteinase K and incubated 1 hour at 50°C. The RNA is extracted using 500µL of Tri reagent (MRC), resuspended in 25µL of water. Equal volumes of samples are reverse transcribed and quantified by quantitative PCR.

## DISCUSSION

In our study we investigated for the first time the CLUH interactome using two complementary proteomic approaches to identify stable and transient CLUH interactions in mammalian cells. We identified both SPAG5 and KNSTRN, also known as Astrin/Kinastrin complex [27], as stable interactants of CLUH in both HCT116 cells and mESC. We showed that this RNA-independent interaction requires the TPR domain of CLUH and the formed complex is localized within granular structures similar to previously described SPAG5 localization at centrosomes [28]. SPAG5, initially identified as being required for the maintenance of sister chromatid cohesion and centrosome integrity [28], has been described as a negative regulator of mTORC1 activation [47]. Interestingly, CLUH has also been recently linked to mTORC1 inhibition upon starvation stress and the mitochondrial clustering phenotype occurring in the absence of CLUH was shown to be rescued by treating the cells with rapamycin, a potent mTOR inhibitor [21]. Considering our findings, it is therefore tempting to speculate that the two proteins may be functioning together in a complex to regulate the mTORC1 signaling. However, this interesting correlation remains to be explored in further studies.

Although CLUH was reported to form G3BP1-positive granules upon stress [21], we did not observe any CLUH or CLUH-SPAG localization in stress granules upon starvation nor oxidative stress in our experimental conditions.

Surprisingly CLUH has been reported to bind SPAG5 mRNA [11], which suggest the existence of different functions for CLUH towards *SPAG5* transcript and its protein. Regarding SPAG5 link to microtubules and CLUH being an RNA binding protein, one could point to a role in mRNP formation and transport.

Although we showed a conservation of SPAG5-CLUH stable interaction in mESC cells, much more significantly enriched proteins were identified by co-IP compared to HCT116 cells. This difference is presumably due to lack of competition between the tagged and the endogenous CLUH, owing to the knock in tagging strategy used in mESCs. Most of those additional proteins correspond to RNA binding proteins, likely pulled downs indirectly *via* CLUH bound mRNAs. SPAG5 interaction with CLUH was also confirmed using the BioID *in vivo* proximity labeling approach in HCT116 cells. Intriguingly SPAG5 was not identified using BioID in mESC cells while it was successfully identified by co-IP. It is possible that in mESC the CLUH-SPAG5 formed complex has a different composition, making SPAG5 not accessible to biotinylation by the N-terminal BioID-tagged CLUH.

In this study we also revealed for the first time a CLUH self-interaction in mammalian cells. Like for SPAG5, this interaction is RNA-independent but requires CLUH TPR domain.

Unlike SPAG5-CLUH interaction, CLUH-CLUH interaction is detected only by Co-IP and not by proximity labeling assay. A possible explanation is that CLUH is likely part of a larger multi-CLUH protein complex that would prevent trans-biotinylation. The interaction of CLUH with itself and with SPAG5 may not be direct and could involve other partners. Interestingly, SPAG5 does not seem required for CLUH self-interaction, as knocking down SPAG5 has no impact of CLUH-CLUH pulldown efficiency.

Whilst the BioID approach confirmed CLUH stable interactants, it principally highlighted unexpected transient interactions with mitochondrial proteins encoded by the nuclear genome. Our study describes a close proximity of CLUH to mitochondrial proteins, that we called CPMPs before their import into mitochondria. Our data suggest that this proximity requires the TPR domain of CLUH and occurs probably during or right after their translation. Indeed, using both confocal microscopy and biochemical fractionation, we did not detect any CLUH accumulation inside mitochondria, indicating that this interaction occurs in a transient manner in the cytosol. The lack of predicted MTS in CLUH and the absence of mitochondria encoded proteins in our BioID data confirmed our observations. In *Drosophila*, CLUH ortholog has been described at the mitochondrial surface [20]. This interaction with the OMM may not conserved in mammals as we did not detect any mitochondrial outer membrane protein in both co-IP and proximity labeling data.

Additionally, using an original TurboID-based time course labeling approach [34], we showed the accumulation of CPMPs overtime suggesting the involvement of CLUH in a dynamic biological process. The analysis of the two time points TurboID data allowed us to discriminate between stable and transient CLUH interactants by taking advantage of the variation of enrichment overtime that is inversely correlated to the residency time near CLUH. Although CLUH could be involved in both protein import and translation, we excluded protein import as we did not detect any chaperone or TOM complex component [3] in our proximity labeling data. On the other hand, we identified several ribosomal subunits, suggesting a proximity with the translation machinery. While in *Drosophila* Clueless has been described to bind the ribosome [20], the mammalian protein has been frequently associated to translation [12,21,22] but a direct interaction with the ribosome has never been formally demonstrated. In line with this, our data suggest a proximity with the translation machinery but not a direct interaction with the ribosome. Indeed, we identified only few ribosomal subunits that are roughly localized at the ribosomal surface, thus more accessible for the biotinlylation by CLUH. We also detected CLUH in the polysomal fraction, suggesting that the protein could be associated with mRNAs in translation. Alike previous reports on CLUH functions in mammalian cells, we observed no effect on mitochondrial protein translation in HCT116 cells.

Interestingly, we found that CLUH also binds the mRNAs of identified CPMPs in a TPR dependent manner, as a deletion of this protein-protein interaction domain abolishes the binding to mRNAs. In agreement with this, the TPR domain of *Drosophila* Clueless was also described to facilitate mRNA binding [20]. Although not strictly demonstrated, our data points towards an association of CLUH with mRNA coding for CPMPs and the persistence of this association during translation may explain CLUH-mediated proximity labeling of CPMPs. The localization of biotinylated residues on CPMPs would be very informative to understand in which translation phase CLUH may be associated. Unfortunately, this information is lost during the on-beads digestion step, due to the too strong affinity of biotin for streptavidin beads.

We identified CLUH proximal proteins in mESC before and after spontaneous EB differentiation. Interestingly, we observed two times more biotinylated proteins in differentiated cells compared to non-differentiated cells. CLUH may therefore be binding a different set of proteins upon the differentiation process, indicating an increased CLUH activity of biological significance. In fact, it could reflect the well-known differences in term of mitochondrial activity and morphology between undifferentiated and differentiated cells [19].

Our data also revealed an interesting proximity of CLUH with some cytoskeleton related proteins and to RNA binding proteins. This could suggest that CLUH forms specific mRNPs with messenger RNA coding for mitochondrial proteins, with implication in storage or sub-cellular localization [48, 49]. This idea is consistent with previous reports on CLUH forming particles [21,50,51] and retaining mitochondrial destined mRNAs in the cytosol [22]. We explored the later possibility by assessing the effect of CLUH on mRNA localization near mitochondria. We observed a higher enrichment of some mRNAs coding for CPMPs in the mitochondrial fraction in CLUH knockout mutant compared to wildtype cells. Interestingly, for those mRNAs we also observed an enrichment in polysomes isolated from the mitochondrial fraction, suggesting an increased translation efficiency. However, in our experimental conditions, this localized-translation effect is completely covered by the main cytosolic translation. We hypothesize that CLUH regulation of localized translation may be relevant only in specific stress, physiological or even pathological conditions.

Altogether, our data suggest that CLUH is associated to mitochondrial mRNAs even during translation, thus facilitating its proximity to mitochondrial proteins prior their import to mitochondria. Nevertheless, the function of this proximity remains to be discovered as we did not observe any effect on translation. Based on our data on mRNA enrichment in the mitochondrial fraction, we speculate that CLUH may be a negative regulator of the targeting of some mRNAs toward the mitochondrial surface.

## ACCESSION NUMBERS

The mass spectrometry proteomics data have been deposited to the ProteomeXchange Consortium via the PRIDE [52] partner repository with the dataset identifiers PXD 027158 and PXD027122.

## SUPPLEMENTARY MATERIAL

Supplementary figures S1-S9

Supplementary figure legends S1-S9

Tables S1 – HCT116 Co-IP spectral count data

Tables S2 – mESC Co-IP spectral count data

Tables S3 – HCT116 BioID spectral count data

Tables S4 – mESC-ES BioID spectral count data

Tables S5 – mESC-EB BioID spectral count data

Tables S6 – HCT116-mito proteome spectral count data

Tables S7 – HCT116 TurboID spectral count data

Tables S8 – Plasmids used in the study

Tables S9 – Oligonucleotides used in the study

Tables S10 – Antibodies used in the study

## ACKNOWLEDGMENTS

We are thankful to E. Girardi, for helpful discussions and critical reading of the manuscript. We acknowledge Philippe Hammann and Lauriane Kuhn from the Plateforme Protéomique Strasbourg Esplanade (CNRS) for their help inperforming the mass-spectrometry analysis.

This project was supported by a grant from French Agence Nationale de la Recherche” ANR (ANR-18-CE12-0021-01 “Polyglot”), by Université de Strasbourg and by the French National Program “Investissement d’Avenir” (Labex MitoCross). M.H was supported by a fellowship from the French Ministère de l’Enseignement Supérieur et de la Recherche. The authors declare no financial or non-financial competing interests.

## AUTHOR CONTRIBUTIONS

RPN conceived, designed and supervised the study. HM, AH and RPN performed the experiments. MH, RPN and AMD analyzed and interpreted the data. RPN wrote the manuscript with inputs from MH and AMD. JC performed MS analysis. AMD provided expertise and secured funding.

## REFERENCES

1. Friedman JR, Nunnari J. Mitochondrial form and function. Nature. 2014;505:335–43.

2. Rath S, Sharma R, Gupta R, Ast T, Chan C, Durham TJ, et al. MitoCarta3.0: an updated mitochondrial proteome now with sub-organelle localization and pathway annotations. Nucleic Acids Res. 2021;49:D1541–7.

3. Bykov YS, Rapaport D, Herrmann JM, Schuldiner M. Cytosolic Events in the Biogenesis of Mitochondrial Proteins. Trends Biochem Sci. The Authors; 2020;:1–18.

4. Béthune J, Jansen R-P, Feldbrügge M, Zarnack K. Membrane-Associated RNA-Binding Proteins Orchestrate Organelle-Coupled Translation. Trends in Cell Biology. 2019;29:178–88.

5. Lesnik C, Golani-Armon A, Arava Y. Localized translation near the mitochondrial outer membrane: An update. RNA Biol. 2015;12:801–9.

6. Fazal FM, Han S, Parker KR, Kaewsapsak P, Xu J, Boettiger AN, et al. Atlas of Subcellular RNA Localization Revealed by APEX-Seq. Cell. 2019;178:473–490.e26.

7. Vincent T, Vingadassalon A, Ubrig E, Azeredo K, Srour O, Cognat V, et al. A genome-scale analysis of mRNAs targeting to plant mitochondria: upstream AUGs in 5’ untranslated regions reduce mitochondrial association. Plant J. Wiley/Blackwell (10.1111); 2017;92:1132– 42.

8. Gehrke S, Wu Z, Klinkenberg M, Sun Y, Auburger G, Guo S, et al. PINK1 and Parkin control localized translation of respiratory chain component mRNAs on mitochondria outer membrane. Cell Metabolism. 2015;21:95–108.

9. Hentze MW, Castello A, Schwarzl T, Preiss T. A brave new world of RNA-binding proteins. Nature Publishing Group. Nature Publishing Group; 2018;19:1–15.

10. Schatton D, Rugarli EI. A concert of RNA-binding proteins coordinates mitochondrial function. Critical Reviews in Biochemistry and Molecular Biology. 2018;53:652–66.

11. Gao J, Schatton D, Martinelli P, Hansen H, Pla-Martin D, Barth E, et al. CLUH regulates mitochondrial biogenesis by binding mRNAs of nuclear-encoded mitochondrial proteins. J Cell Biol. 2014;207:213–23.

12. Schatton D, Pla-Martin D, Marx M-C, Hansen H, Mourier A, Nemazanyy I, et al. CLUH regulates mitochondrial metabolism by controlling translation and decay of target mRNAs. J Cell Biol. 2017;216:675–93.

13. Logan DC, Scott I, Tobin AK. The genetic control of plant mitochondrial morphology and dynamics. The Plant Journal. 2003;36:500–9.

14. Cox RT, Spradling AC. Clueless, a conserved Drosophila gene required for mitochondrial subcellular localization, interacts genetically with parkin. Disease Models & Mechanisms. 2009;2:490–9.

15. Fields SD, Arana Q, Heuser J, Clarke M. Mitochondrial membrane dynamics are altered in cluA-mutants of Dictyostelium. J. Muscle Res. Cell. Motil. 2002;23:829–38.

16. Fields SD, Conrad MN, Clarke M. The S. cerevisiae CLU1 and D. discoideum cluA genes are functional homologues that influence mitochondrial morphology and distribution. J. Cell. Sci. The Company of Biologists Ltd; 1998;111:1717–27.

17. Wakim J, Goudenege D, Perrot R, Gueguen N, Desquiret-Dumas V, Chao de la Barca JM, et al. CLUH couples mitochondrial distribution to the energetic and metabolic status. J. Cell. Sci. 2017;130:1940–51.

18. Ellen Kreipke R, Wang Y, Miklas JW, Mathieu J, Ruohola-Baker H. Metabolic remodeling in early development and cardiomyocyte maturation. Semin. Cell Dev. Biol. 2016;52:84–92.

19. Lisowski P, Kannan P, Mlody B, Prigione A. Mitochondria and the dynamic control of stem cell homeostasis. EMBO Rep. EMBO Press; 2018;19:e45432–12.

20. Sen A, Cox RT. Clueless is a conserved ribonucleoprotein that binds the ribosome at the mitochondrial outer membrane. Biology Open. 2016;5:195–203.

21. Pla-Martin D, Schatton D, Wiederstein JL, Marx M-C, Khiati S, Krüger M, et al. CLUH granules coordinate translation of mitochondrial proteins with mTORC1 signaling and mitophagy. EMBO J. 2020;39:1717–23.

22. Vardi-Oknin D, Arava Y. Characterization of Factors Involved in Localized Translation Near Mitochondria by Ribosome-Proximity Labeling. Front Cell Dev Biol. 2019;7:305.

23. Kamiyama D, Sekine S, Barsi-Rhyne B, Hu J, Chen B, Gilbert LA, et al. Versatile protein tagging in cells with split fluorescent protein. Nature Communications. Nature Publishing Group; 2016;7:11046–9.

24. Kim DI, Jensen SC, Noble KA, Kc B, Roux KH, Motamedchaboki K, et al. An improved smaller biotin ligase for BioID proximity labeling. Zheng Y, editor. Molecular Biology of the Cell. American Society for Cell Biology; 2016;27:1188–96.

25. Branon TC, Bosch JA, Sanchez AD, Udeshi ND, Svinkina T, Carr SA, et al. Efficient proximity labeling in living cells and organisms with TurboID. Nat Biotechnol. Nature Publishing Group; 2018;36:1–23.

26. Teo G, Liu G, Zhang J, Nesvizhskii AI, Gingras A-C, Choi H. SAINTexpress: improvements and additional features in Significance Analysis of INTeractome software. Journal of Proteomics. 2014;100:37–43.

27. Dunsch AK, Linnane E, Barr FA, Gruneberg U. The astrin-kinastrin/SKAP complex localizes to microtubule plus ends and facilitates chromosome alignment. The Journal of Cell Biology. 2011;192:959–68.

28. Thein KH, Kleylein-Sohn J, Nigg EA, Gruneberg U. Astrin is required for the maintenance of sister chromatid cohesion and centrosome integrity. J Cell Biol. 2007;178:345–54.

29. Zhong W, Zhou Y, Li J, Mysore R, Luo W, Li S, et al. OSBP-related protein 8 (ORP8) interacts with Homo sapiens sperm associated antigen 5 (SPAG5) and mediates oxysterol interference of HepG2 cell cycle. Experimental Cell Research. Elsevier; 2014;322:227–35.

30. Blatch GL, Lässle M. The tetratricopeptide repeat: a structural motif mediating protein-protein interactions. BioEssays. John Wiley & Sons, Ltd; 1999;21:932–9.

31. Qin W, Cho KF, Cavanagh PE, Ting AY. Deciphering molecular interactions by proximity labeling. Nat Meth. Springer US; 2020;116:1–11.

32. Boos F, Labbadia J, Herrmann JM. How the Mitoprotein-Induced Stress Response Safeguards the Cytosol: A Unified View. Trends in Cell Biology. Elsevier Ltd; 2020;30:1–14.

33. Wrobel L, Topf U, Bragoszewski P, Wiese S, Sztolsztener ME, Oeljeklaus S, et al. Mistargeted mitochondrial proteins activate a proteostatic response in the cytosol. Nature. Nature Publishing Group; 2015;524:485–8.

34. Cho KF, Branon TC, Udeshi ND, Myers SA, Carr SA, Ting AY. Proximity labeling in mammalian cells with TurboID and split-TurboID. Nat Protoc. Springer US; 2020;15:1–31.

35. Ngondo RP, Cirera-Salinas D, Yu J, Wischnewski H, Bodak M, Vandormael-Pournin S, et al. Argonaute 2 Is Required for Extra-embryonic Endoderm Differentiation of Mouse Embryonic Stem Cells. Stem Cell Reports. ElsevierCompany; 2018;10:461–76.

36. Ngondo RP, Cohen-Tannoudji M, Ciaudo C. Fast In Vitro Procedure to Identify Extraembryonic Differentiation Defect of Mouse Embryonic Stem Cells. STAR Protocols. The Author(s); 2020;1:100127.

37. Gibson DG, Young L, Chuang R-Y, Venter JC, Hutchison CA, Smith HO. Enzymatic assembly of DNA molecules up to several hundred kilobases. Nat Meth. Nature Publishing Group; 2009;6:343–5.

38. Schindelin J, Arganda-Carreras I, Frise E, Kaynig V, Longair M, Pietzsch T, et al. Fiji: an open-source platform for biological-image analysis. Nat Meth. Nature Publishing Group; 2012;9:676–82.

39. Klickstein JA, Mukkavalli S, Raman M. AggreCount: an unbiased image analysis tool for identifying and quantifying cellular aggregates in a spatially defined manner. J Biol Chem. 2020;295:17672–83.

40. Mutterer J, Zinck E. Quick-and-clean article figures with FigureJ. J Microsc. John Wiley & Sons, Ltd; 2013;252:89–91.

41. Lange H, Ndecky SYA, Gomez-Diaz C, Pflieger D, Butel N, Zumsteg J, et al. RST1 and RIPR connect the cytosolic RNA exosome to the Ski complex in Arabidopsis. Nature Communications. 2019;10:3871.

42. Bouyssié D, Hesse A-M, Mouton-Barbosa E, Rompais M, Macron C, Carapito C, et al. Proline: an efficient and user-friendly software suite for large-scale proteomics. Bioinformatics (Oxford, England). 2020;36:3148–55.

43. Raudvere U, Kolberg L, Kuzmin I, Arak T, Adler P, Peterson H, et al. g:Profiler: a web server for functional enrichment analysis and conversions of gene lists (2019 update). Nucleic Acids Res. 2019;47:W191–8.

44. Doncheva NT, Morris JH, Gorodkin J, Jensen LJ. Cytoscape StringApp: Network Analysis and Visualization of Proteomics Data. J. Proteome Res. American Chemical Society; 2019;18:623–32.

45. Williamson CD, Wong DS, Bozidis P, Zhang A, Colberg-Poley AM. Isolation of Endoplasmic Reticulum, Mitochondria, and Mitochondria-Associated Membrane and Detergent Resistant Membrane Fractions from Transfected Cells and from Human Cytomegalovirus-Infected Primary Fibroblasts. Current Protocols in Cell Biology. Hoboken, NJ, USA: John Wiley & Sons, Inc; 2001. pp. 3.27.1–3.27.33.

46. Pringle ES, McCormick C, Cheng Z. Polysome Profiling Analysis of mRNA and Associated Proteins Engaged in Translation. Curr Protoc Mol Biol. John Wiley & Sons, Ltd; 2019;125:e79.

47. Thedieck K, Holzwarth B, Prentzell MT, Boehlke C, Kläsener K, Ruf S, et al. Inhibition of mTORC1 by Astrin and Stress Granules Prevents Apoptosis in Cancer Cells. Cell. 2013;154:859–74.

48. Mitchell SF, Parker R. Principles and Properties of Eukaryotic mRNPs. Mol. Cell. Elsevier Inc; 2014;54:547–58.

49. Hentze MW, Castello A, Schwarzl T, Preiss T. A brave new world of RNA-binding proteins. Nature Publishing Group. Nature Publishing Group; 2018;19:327–41.

50. Sheard KM, Thibault-Sennett SA, Sen A, Shewmaker F, Cox RT. Clueless forms dynamic, insulin-responsive bliss particles sensitive to stress. Developmental Biology. Elsevier Ltd; 2020;459:149–60.

51. Ando T, Yamayoshi S, Tomita Y, Watanabe S, Watanabe T, Kawaoka Y. The host protein CLUH participates in the subnuclear transport of influenza virus ribonucleoprotein complexes. Nat Microbiol. 2016;1:16062.

52. Perez-Riverol Y, Csordas A, Bai J, Bernal-Llinares M, Hewapathirana S, Kundu DJ, et al. The PRIDE database and related tools and resources in 2019: improving support for quantification data. Nucleic Acids Res. 2019;47:D442–50.

