## Supplementary figures and images for "CLUH interactome reveals an association to SPAG5 and a proximity to the translation of mitochondrial protein"

### Supplemental Figures

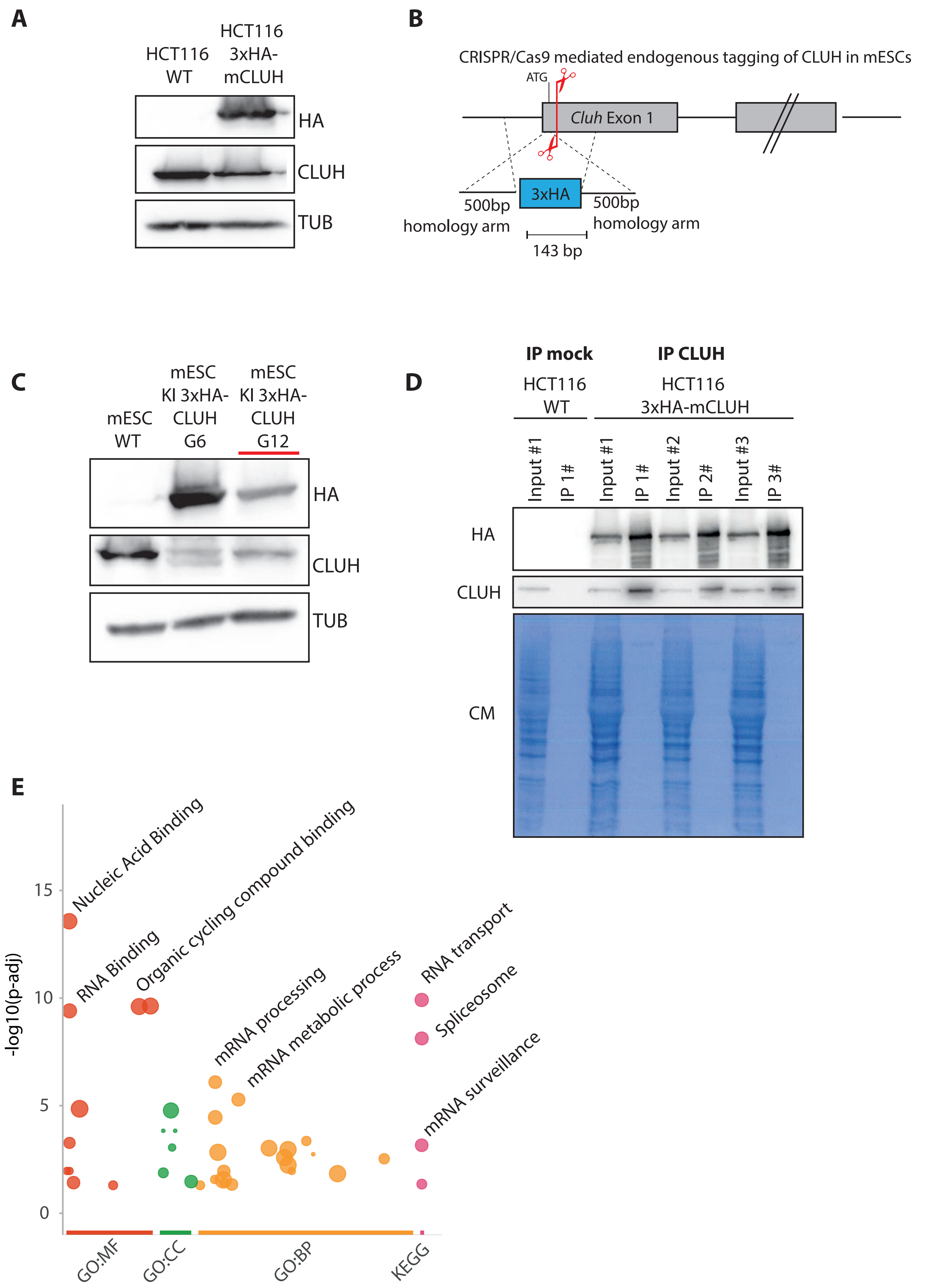

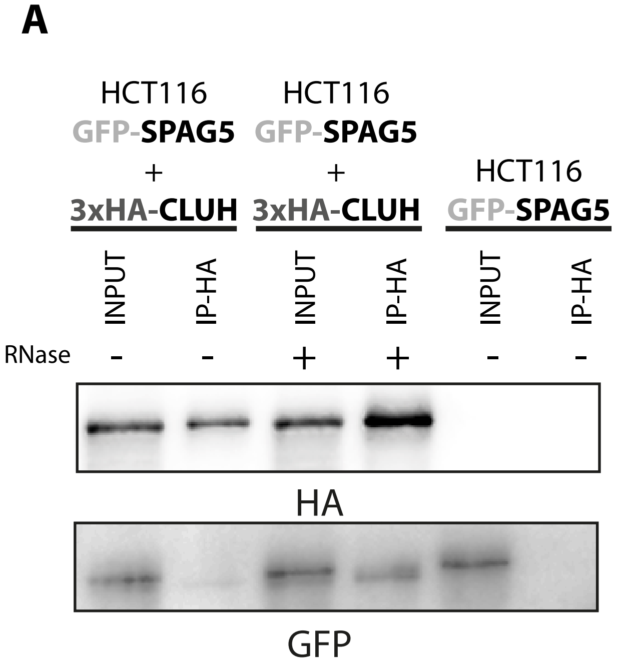

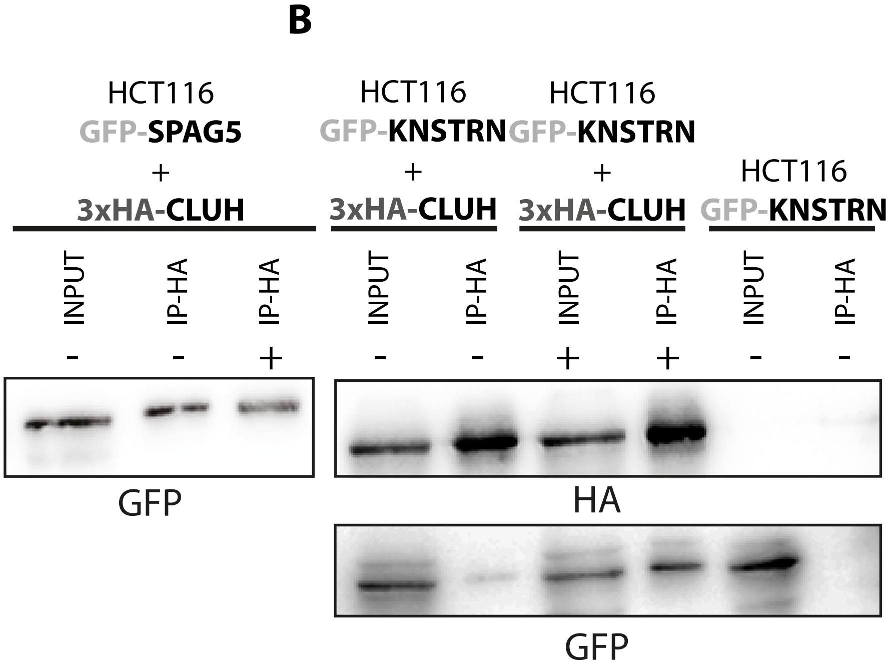

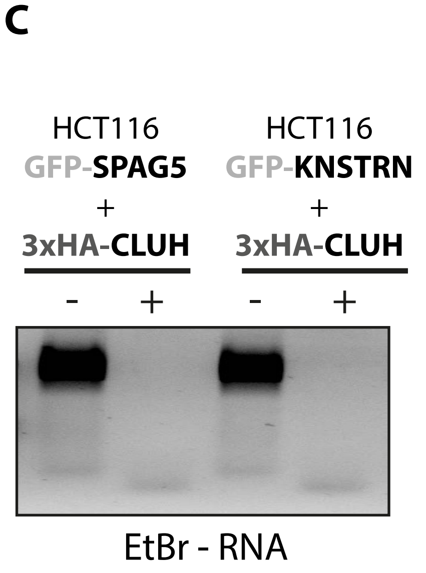

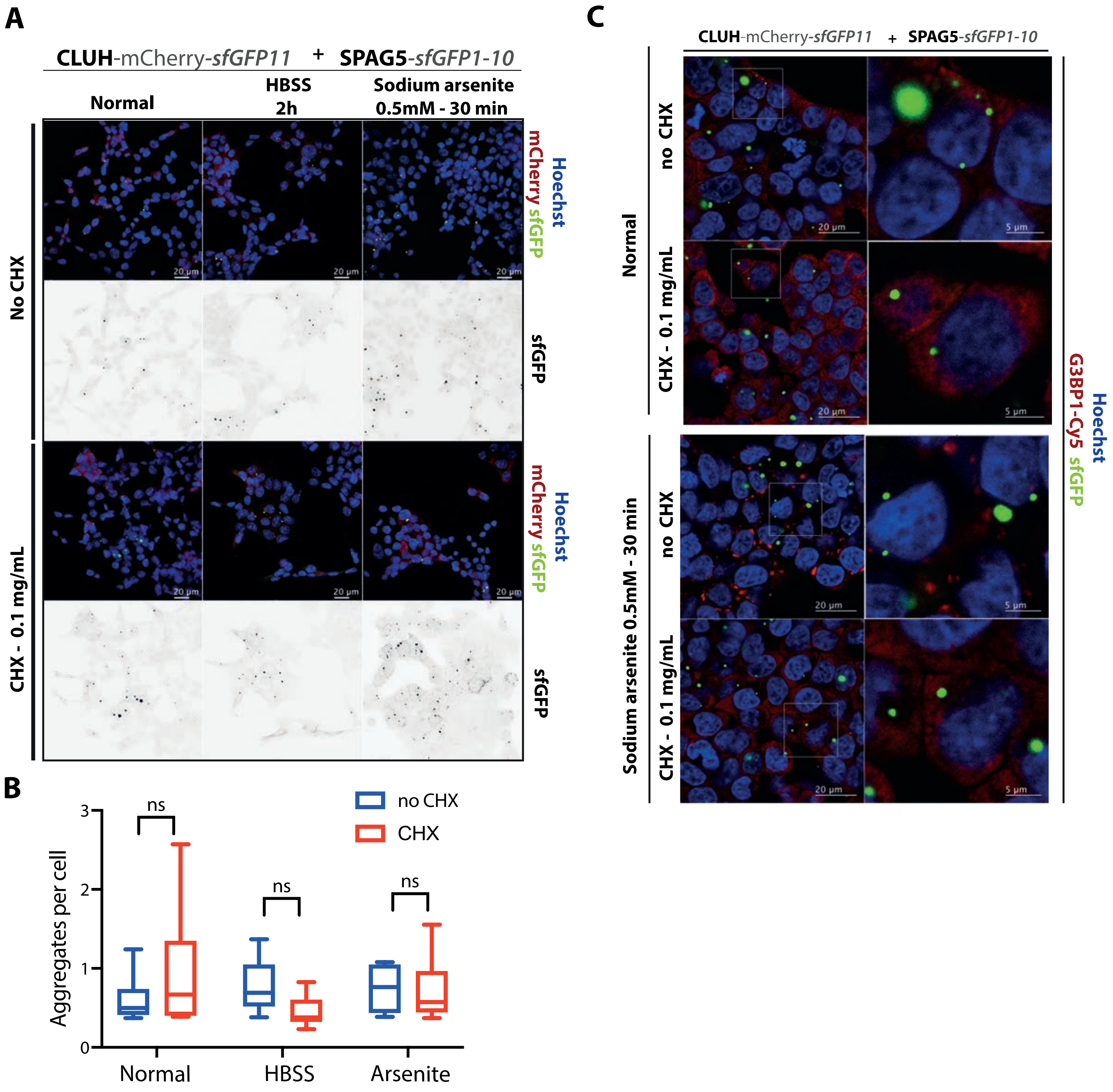

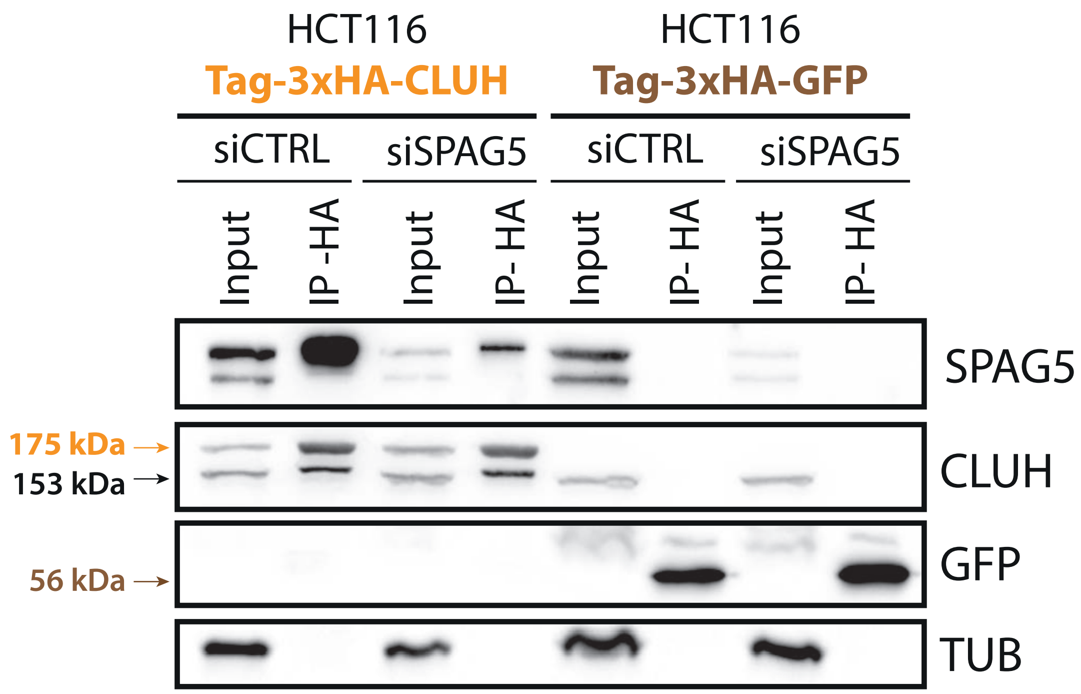

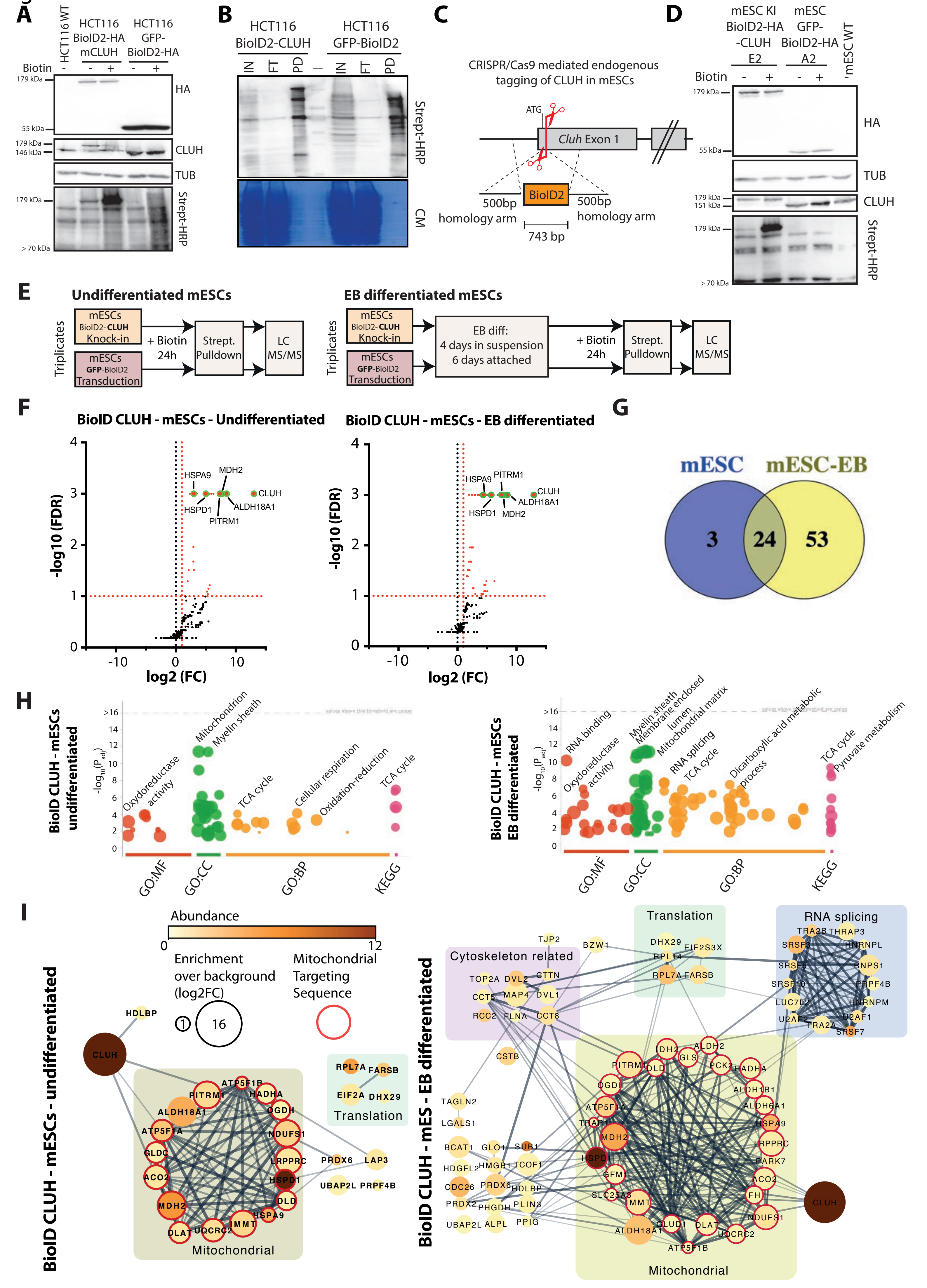

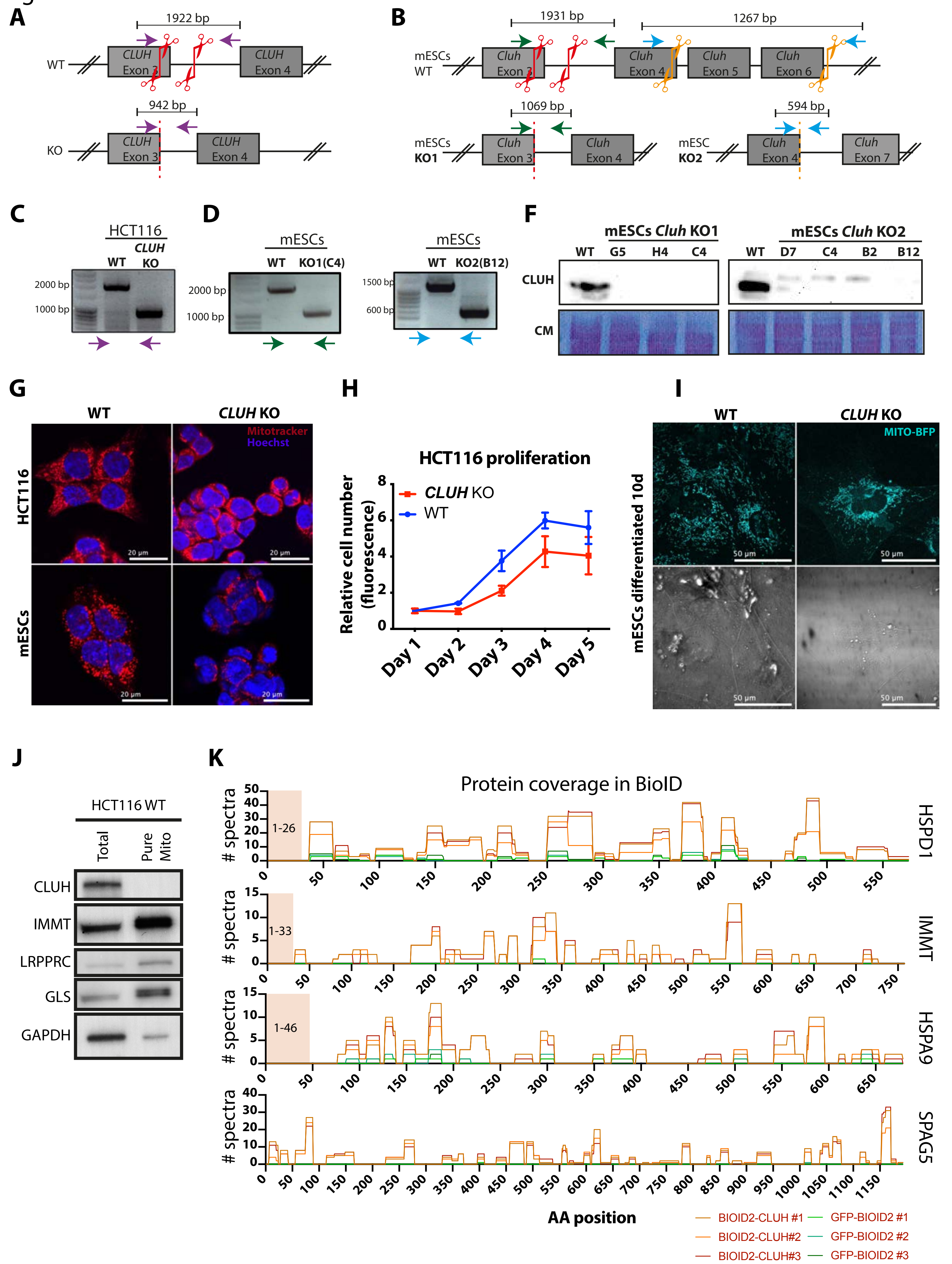

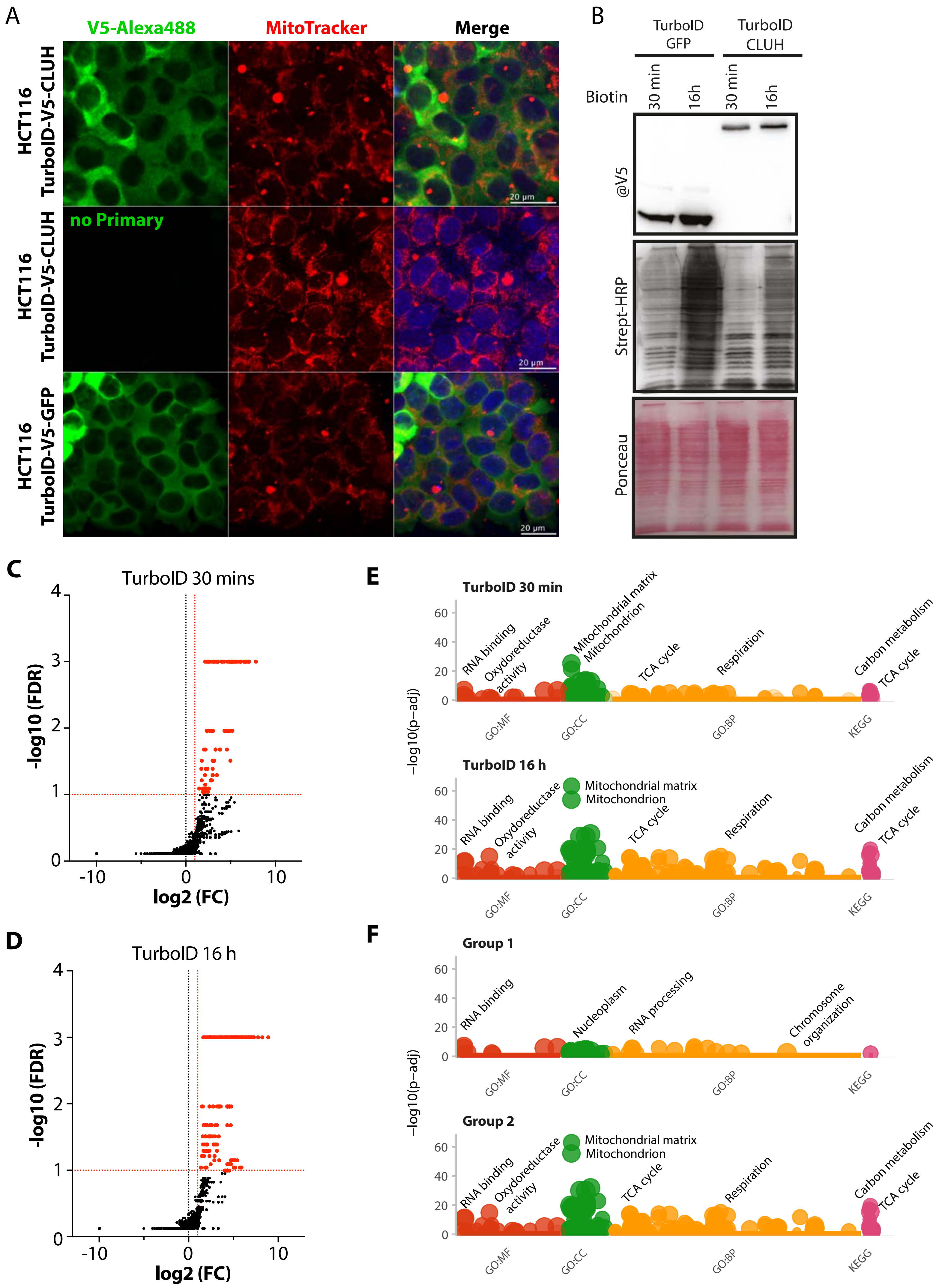

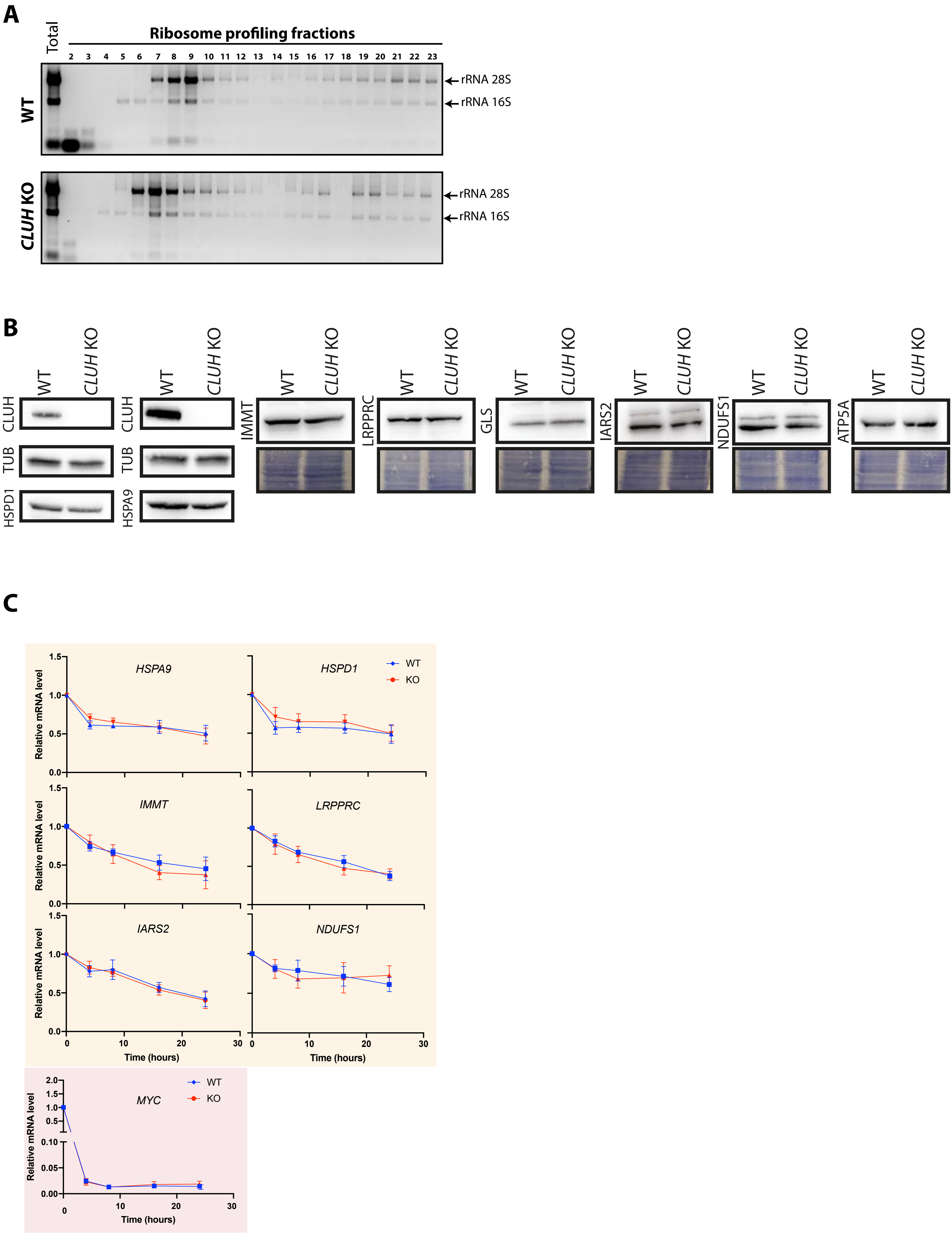

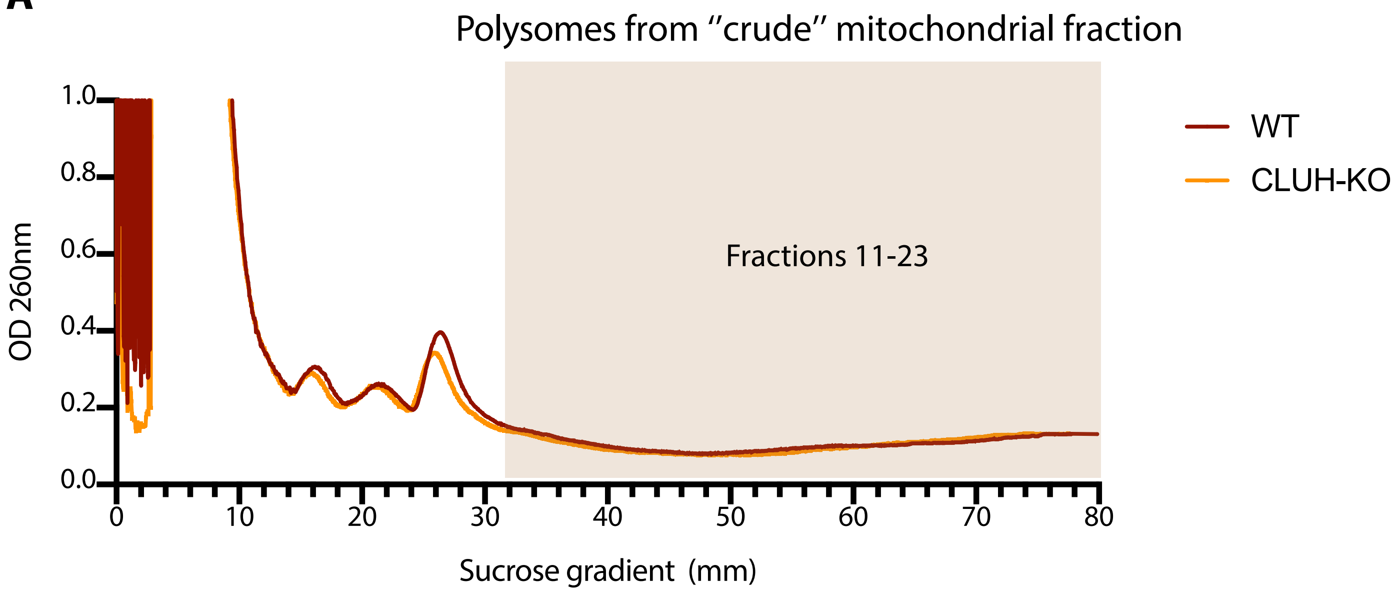

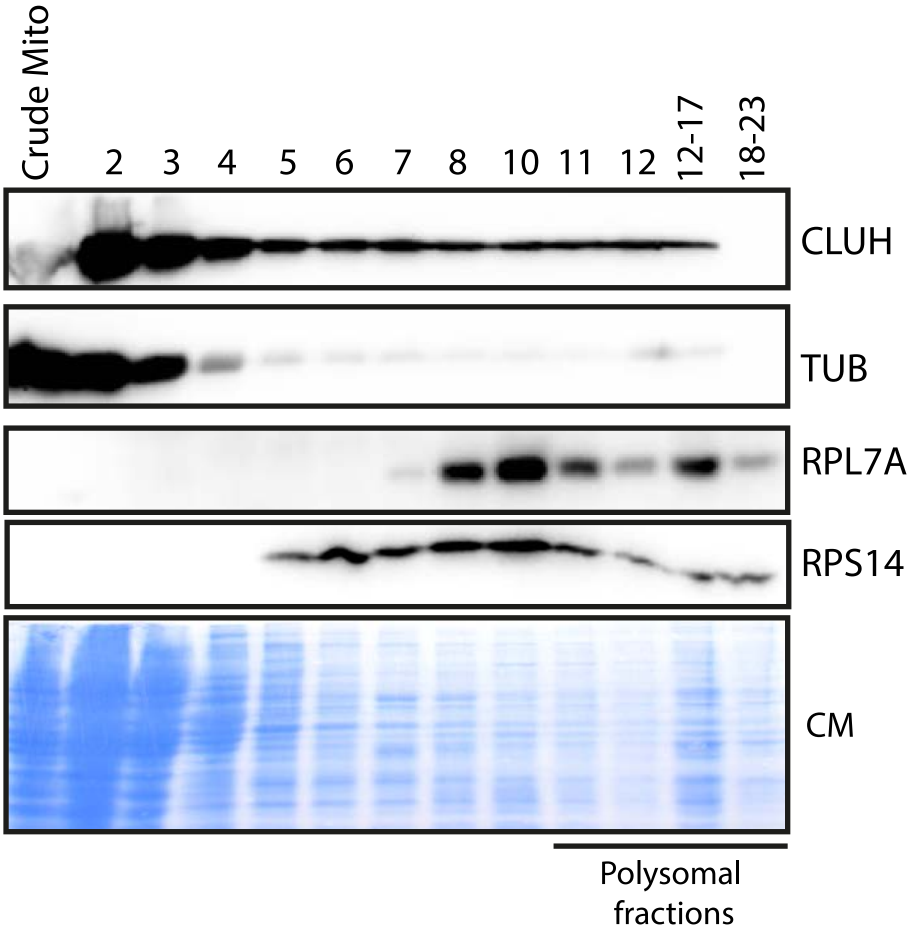
