## Supplemental Figure legends for "CLUH interactome reveals an association to SPAG5 and a proximity to the translation of mitochondrial protein"

**Figure S1: Generation of 3xHA-CLUH expressing cells and proteomic analysis. (A)** Western blot showing the expression of N-terminal HA-tagged mouse CLUH protein (mCLUH) in polyclonal HCT116 stable cell line. Indicated proteins are detected using specific antibodies. **(B)** Schematic representation of the CRISPR/Cas9 mediated knock-in strategy in mESCs, to endogenously tag CLUH in N-terminal with 3xHA. Red scissors indicate the cleavage site**. (C)** Immunodetection analysis of the endogenously 3xHA-tagged CLUH protein in mESCs. Two selected clones (G6 and G12) are analyzed. The proteins are detected using antibodies directed against CLUH and the HA tag. Coomassie (CM) staining of the membrane is used as loading control. The clone underlined in red was used for the co-IP experiment. **(D)** Western blot analysis of a fraction of total extracts and Co-IP replicate samples sent for LC-MS/MS. One representative control (HCT116 WT) experiment is shown. The 3xHA-mCLUH protein is detected using anti-HA antibodies, CLUH is detected using specific antibodies and the Coomassie (CM) staining of the membrane is used as loading control. **(E)** Manhattan plot illustrating the gene ontology and pathway enrichment analysis of proteins identified in mESC CLUH co-IP experiment, generated using G:profiler tool [43]. The functional terms, associated with the protein list, are grouped in four categories: GO: MF (Molecular Function), GO : BP (Biological Process), GO : CC (Cellular Component) and KEGG pathways. The y-axis shows the adjusted enrichment p-values in negative log10 scale. The circle sizes are in accordance with the corresponding term size (in the data base) and terms from the same GO subtree are located close to each other on the x-axis. The most significantly enriched terms are indicated

**Figure S2: RNA-independent CLUH interaction with SPAG5 and KNSTRN.**

**(A-B)** Western blot analysis of co-IP, between 3xHA-CLUH and GFP-SPAG5 (**A**) or GFP-tagged KNSTRN (**B**) stably expressed in HCT116 cells. The co-IP is performed using magnetic beads coupled with anti-HA antibodies (IP-HA) on total extracts (INPUT) treated (+) or not (-) with RNaseA/T1. Proteins are detected using anti-HA and anti-GFP antibodies. **(C)** Ethidium bromide-stained agarose gel loaded with RNase treated (+) or non-treated (-) total protein extracts used for the IP showing the presence of ribosomal RNA.

**Figure S3: CLUH-SPAG5 formed structures are not stress granules.**

**(A)** Split-GFP analysis of CLUH and SPAG5 interaction on fixed cells expressing both CLUH fused with mCherry-sfGFP11 and SPAG5 fused with sfGFP1-10. The constructs are stably expressed in HCT116 cells cultured in standard conditions (normal), nutritional stress medium for 2 hours (HBSS, low-glucose medium devoid of serum and amino acids) and oxidative stress condition for 30 minutes (0.5 mM sodium arsenite). Treatment with 0.1 mg/mL of cycloheximide (CHX) or control (No CHX) is shown. The reconstituted sfGFP signal is shown in green (upper panels) and in black (lower panels). mCherry signal is shown in red and nuclei, stained with Hoechst, are in blue. **(B)** Quantification of the sfGFP signal aggregates detected with (red) or without (blue) CHX treatment in normal, HBSS and sodium arsenite conditions. Box plots represent the mean aggregate count per cells from 6 different images containing from 40 to 200 cells each. The quantification is performed using the ImageJ AggreCount macro [39]. The statistical non significance (ns) of the difference due to CHX treatment is determined using a t-test with a threshold p-value of 0.05. **(C)** Immunofluorescence analysis of Split-GFP experiment on fixed HCT116 cells stably expressing both CLUH-mCherry-sfGFP11 and SPAG5-sfGFP1-10 constructs. The cells cultured in standard conditions (normal) and oxidative stress conditions (Sodium arsenite) for 30 minutes are treated or not with CHX. The signal of marker G3BP1 is revealed using a specific antibody and secondary antibodies coupled with Cy5 and shown in red. The mCherry signal in not shown. The reconstituted sfGFP signal is shown in green and nuclei, stained with Hoechst, are in blue.

**Figure S4: SPAG5 is not required for CLUH-self interaction.**

Western blot analysis of CLUH self-interaction and its interaction with SPAG5 by co-IP. HCT116 cells stably expressing BioID2-3xHA-CLUH (Tag-3xHA-CLUH) protein are transfected with siRNA directed against SPAG5 (siSPAG5) or with non-targeting control siRNA (siCTRL). HCT116 expressing identically tagged GFP protein (Tag-3xHA-GFP) is used as a control. The co-IP is performed on total protein extracts (INPUT) using magnetic beads couplet with anti-HA antibodies (IP-HA). The indicated proteins are revealed using specific antibodies. The molecular weight of the tagged CLUH (orange), the endogenous CLUH (black) and GFP (brown) is indicated. TUBULIN (TUB) is used as a loading control.

**Figure S5: Identification of CLUH proximal proteins using BioID in undifferentiated and EB differentiated mESCs.**

Western blot analysis of the expression of BioID2-HA-CLUH and GFP-BioID2-HA proteins in HCT116 derived polyclonal stable cell line. Total extract from wild-type HCT116 cells (HCT116 WT) is used a control. CLUH and Tubulin (TUB) are revealed using specific antibodies. The cells were incubated in the presence of 50µM biotin for 24h and global biotinylation activity revealed using HRP-coupled streptavidin (Strept-HRP). **(B)** Representative image of the western blot analysis of one replicate BioID experiment performed on HCT116 cells and send for LC-MS/MS analysis. Equal volumes of both the input protein extract (IN) and the flow through (FT) as well as a 1/10 fraction of the pulldown (PD) sample are analyzed. The biotinylated proteins are revealed using HRP-coupled streptavidin (Strept-HRP). The

Coomassie staining of the membrane (CM) is used as loading control. **(C)** Schematic representation of the CRISPR/Cas9 mediated knock-in strategy in mESCs to endogenously tag CLUH in N-terminal with BioID2-HA. Red scissors indicate the cleavage site **(D)** Western blot analysis of the expression of BioID2-HA-CLUH in CRISPR/Cas9 generated E2 knock-in clone and of the expression of GFP-BioID2 protein in mESC (E14) derived stable cell line. Total extract from wild-type ESCs (mESCs WT) is used a control. CLUH and Tubulin (TUB) are revealed using specific antibodies. The cells were incubated in the presence of 50µM biotin for 24h and global biotinylation activity revealed using HRP-coupled streptavidin. **(E)** Schematic representation of the BioID experimental design using undifferentiated (left) and embryoïd bodies (EB) differentiated (right) mESCs expressing the BioID2 protein fused to CLUH (endogenous tagging, clone E2) or to GFP (stable cell line generated by lentiviral transduction) proteins. The proximity labeling is performed for 24 hours in the presence of 50 µM biotin in the medium. Biotinylated proteins, from both the specific (BioID2-CLUH) and control (GFP-BioID2) samples, are isolated using streptavidin-coupled magnetic beads and identified by Liquid Chromatography coupled to tandem Mass Spectrometry (LC MS/MS). **(F)** Volcano plots showing the global enrichment of proteins in BioID2-CLUH versus the GFP-BioID2 control in both undifferentiated (left) and EB differentiated (right) mESC. The x-axis shows the log2 fold change (FC), and the y-axis shows the −log10 of the false discovery rate (n=3), obtained using SAINTexpress software [26]. Significantly enriched proteins are shown in red and are defined by a fold change greater than two and a FDR < 0.1 (shown as dashed red lines). CLUH and five of the most abundant mitochondrial proteins are labeled and identified with a green circle. **(G)** Venn diagram showing the intersection of CLUH proximal proteins, identified by BioID, in undifferentiated (mESCs) and E EB differentiated cells (EB). **(H)** Manhattan plot illustrating the gene ontology and pathway enrichment analysis of proteins identified in BioID experiment on undifferentiated (left) and differentiated (right) mESCs, generated using g:profiler tool [43]. The functional terms, associated with the protein lists, are grouped in four categories: GO: MF (Molecular Function), GO : CC (Cellular Component), GO : BP (Biological Process) and KEGG pathways. The y-axis shows the adjusted enrichment p-values in negative log10 scale. The circle sizes are in accordance with the corresponding term size (in the database) and terms from the same GO subtree are located close to each other on the x-axis. The more significantly enriched terms are labeled. **(I)** Visualization of the functional interaction network of CLUH proximal proteins identified by BioID on undifferentiated (left) and EB differentiated (right) mESCs, generated using the Cytoscape StringApp [44]. The proteins have been grouped according to the most represented functional categories: “Cytoskeleton related”, “Translation”, “Mitochondrial” and “RNA splicing”. The confidence score of each interaction is mapped to the edge thickness and opacity. The size of the node relates to the enrichment in log2 fold change (log2FC) over the BioID-GFP background control. The protein abundance in the BioID2-CLUH sample is illustrated by the color scale and corresponds to the specific spectral count normalized to the protein size. Proteins with mitochondrial targeting sequences (MTS) according to Uniprot database are highlighted in red.

**Figure S6: Generation of CRISPR/Cas9 knockout cells for *CLUH* and BioID protein coverage.**

**(A-B)** Schematic representation of the CRISPR/Cas9 mediated knock-out of *CLUH* in HCT116 **(A)** and mESCs **(B).** A paired sgRNA strategy is used to delete DNA fragments leading to gene inactivation. Red scissors indicate the cleavage site of each sgRNA and colored arrows show the location of PCR genotyping primers. PCR amplification products size are indicated for both wild-type cells and each mutant. **(C-D)** Agarose gel showing the genotyping PCR results of WT and mutant HCT116 **(C)** and mESCs **(D)**. The used PCR primers match the color code indicated in **(A, B).** **(F)** Western blot showing the expression of *Cluh* in selected mESC *Cluh* KO clones. CLUH is detected using specific antibodies. Coomassie staining of the membrane (CM) is used as loading control. **(G)** Confocal microscopy images of mESC and HCT116 *CLUH* KO cells. Parental wildtype cells are also shown. The mESC *Cluh* KO1 (C4) clone is shown. Mitochondria (red) are labeled using MitoTracker™ Red CMXRos. Nuclei (blue) are stained with Hoechst. The scale bar is indicated in white. **(H)** Proliferation assay performed on HCT116 *CLUH* KO cells and WT cells over 5 days. The relative cell number is measured compared to day1, using a florescence assay (CellTiter-FluorTM, Promega). The error bars correspond to the standard deviation of three biological replicate experiments. **(I)** Confocal microscopy images of both WT and *Cluh* KO (C4) mESCs stably expressing mitochondrial BFP protein (fusion with Cox8a mitochondrial targeting sequence). Mitochondria are shown in light blue (upper panels) and transmitted light images are shown in gray. The scale bar is indicated in white. **(J)** Western blot on total and "pure" mitochondrial fractions from wild-type HCT116 cells. Indicated proteins are revealed using specific antibodies. **(K)** Plot showing the protein coverage from LC MS/MS identification of the most abundant proteins from CLUH BioID experiment on HCT116 cells (Figure 4). The x-axis corresponds to the amino acid (aa) position for each protein and the y-axis show specific spectral counts for both BioID2-CLUH and GFP-BIOID2 samples. All replicate samples are indicated with a color scale. The N-terminal region of the proteins, containing the mitochondrial targeting peptide, according to uniport annotatios, is highlighted in orange.

**Figure S7: Identification of CLUH proximal proteins in HCT116 cells using a TurboID time course approach.**

**(A)** Confocal microscopy images of HCT116 cells stably expressing TurboID-V5-CLUH or TurboID-V5-GFP fusion proteins. The proteins are detected using anti-V5 primary antibody and revealed using Alexa488 secondary antibody (green). Mitochondria (red) are labeled using MitoTracker™ Red CMXRos. Nuclei (blue) are stained with Hoechst. The scale bar is indicated in white. **(B)** Western blot showing the expression of TurboID-GFP and TurboID-CLUH constructs stably expressed in HCT116 cells, at 30 min and 16h after the addition of biotin in the medium (Figure 5A). The proteins are revealed using anti-V5 antibodies. Biotinylated proteins are revealed using HRP-coupled streptavidin. Ponceau staining of the membrane is shown as loading control. **(C-D)** Volcano plots showing the global enrichment of proteins in TurboID-CLUH versus the TurboID-GFP control at 30 minutes **(C)** and 16 hours **(D)**. The x-axis shows the log2 fold change (FC), and the y-axis shows the −log10 of the false discovery rate (n=3), obtained using SAINTexpress software [26]. Significantly enriched proteins are shown in red and are defined by a fold change greater than two and a FDR < 0.1 (shown as dashed red lines). **(E-F)** Manhattan plots illustrating the gene ontology and pathway enrichment analysis of proteins identified in TurboID experiment. The analysis is done on all significantly enriched proteins identified at 30 minutes and 16 hours **(E)** as well as for the proteins from group 1 and group 2 **(F)** from figure 5B). The plot is generated using g:profiler tool [43]. The functional terms, associated with the protein lists, are grouped in four categories: GO: MF (Molecular Function), GO : CC (Cellular Component), GO : BP (Biological Process) and KEGG pathways. The y-axis shows the adjusted enrichment p-values in negative log10 scale. The circle sizes are in accordance with the corresponding term size (in the data base) and terms from the same GO subtree are located close to each other on the x-axis. The more significantly enriched terms are labeled.

**Figure S8: CLUH effect on RNA stability and translation.**

**(A)** Representative agarose gel electrophoresis analysis of the RNA extracted from the ribosome profiling fractions of both *CLUH* KO and HCT116 cells. The fractions corresponding to about 3.3 mm of the sucrose gradient and are numbered from 1 to 23. Input RNA from total extract (Total) is loaded as control. 28S and 16S ribosomal RNA are indicated. RNA is revealed using ethidium bromide staining. **(B)** Western blot analysis of mitochondrial proteins abundance in total protein extracts from *CLUH* KO and WT HCT116 cells. Indicated proteins are revealed using specific antibodies. TUBULIN (TUB) or Coomassie staining of the membrane is used as loading control. **(C)** Analysis of mRNA stability over time, after actinomycin D treatment in *CLUH* KO and WT HCT116 cells. The represented mRNA levels are normalized to *GAPDH* levels and are represented relative to time 0. The error bar represents the standard deviation of three replicate experiments. *MYC* is used as a control for the Actinomycin treatment.

**Figure S9: Polysome profiling from crude mitochondrial extract.**

**(A)** Representative graphs of polysome profilings from “crude” mitochondrial fractions (see Figure 7D) of WT and *CLUH* KO HCT116 cells. The y-axis corresponds to the absorbance at 260 nm and the x-axis to the distance in the sucrose gradient. The polysomal fractions used for further experiments are highlighter in orange. **(B)** Representative western blot analysis of polysome profiling from the crude mitochondrial fraction in WT HCT116 cells. Each fraction corresponds to a about 3.3 mm of the sucrose gradient and are numbered from 1 to 23. Fractions 12-17 and 18-23 are pooled. Crude mitochondrial extract is used as control. Indicated proteins are revealed using specific antibodies. Coomassie staining of the membrane is shown as loading control. The polysomal fraction used for RNA extraction are underlined.
